# The Human Hippocampus Contributes to Egocentric Coding of Distance to a Local Landmark

**DOI:** 10.1101/465997

**Authors:** Xiaoli Chen, Paula Vieweg, Thomas Wolbers

## Abstract

Spatial navigation can depend on path integration or environmental cues (e.g., landmarks), which are thought to be integrated in hippocampal and entorhinal circuits. This study investigates the anatomical basis of path integration and navigation based on a single local landmark using an individual differences approach, since people vary substantially in their ability to navigate with path integration cues and landmarks. In two experiments, we dissociated the use of path integration and a local landmark in the same navigation task, and investigated whether morphological variability in the hippocampus and entorhinal cortex could explain behavioral variability in young healthy humans. In Experiment 1, participants navigated in a fully immersive virtual reality environment, with body-based cues available for path integration. The participants first walked through a series of posts before attempting to walk back to the remembered location of the first post. We found that gray matter volume of the hippocampus positively predicted behavioral accuracy of retrieving the target’s distance in relation to the local landmark. Hippocampus also positively predicted path integration performance in terms of walking-distance to the target location. Experiment 2 was conducted in a desktop virtual environment, with no body-based cues available. Optic flow served as path integration cues, and participants were tested on their memory of a learned target location along a linear track. Consistent with Experiment 1, the results showed that hippocampal volume positively predicted performance on the target’s distance in relation to the local landmark. In contrast to Experiment 1, there was no correlation between hippocampal volume and path integration performance. Together, our two experiments provide novel and converging evidence that the hippocampus plays an important role in encoding egocentric distance to a single local landmark during navigation, and they suggest a stronger hippocampal involvement when path integration is based on body-based compared to optic flow cues.

## 1 INTRODUCTION

One cannot survive without the ability to navigate our complex environments, which can be based either on self-motion cues or on environmental cues. Navigating with self-motion cues requires continuous integration of self-motion inputs over distance, a process referred to as path integration (Mittelstaedt & Mittelstaedt, 1980). On the contrary, navigation with environmental cues is a relatively discrete process, since environmental cues are directly informative of the navigator’s position and orientation. During navigation, both cue types typically compensate each other, in that environmental cues can help correct errors accumulated during path integration (Etienne, Maurer, & Seguinot, 1996), while path integration can be used to distinguish between ambiguous environmental cues (Etienne et al., 1996) and to judge the reliability of environmental cues (Zhao & Warren, 2015). Importantly, environmental-cue-navigation and path integration appear to involve distinct cognitive processes (Chen, McNamara, Kelly, & Wolbers, 2017) and differential neural mechanisms (Connor & Knierim, 2017; Knierim, Neunuebel, & Deshmukh, 2014). Given that people differ substantially on the efficiency of utilizing the two cue types (Chen et al., 2017), it is possible to infer neural mechanisms of environmental-cue-navigation and path integration by examining how brain morphology correlates with navigational performance from an individual differences approach.

Previous studies have examined anatomical correlates of different aspects of spatial navigation performance from an individual differences perspective. For example, studies on topographical memory of complex landmark layouts have found that anatomical variability in the hippocampus (HPC) correlates with navigation performance (Bohbot, Lerch, Thorndycraft, Iaria, & Zijdenbos, 2007; Hartley & Harlow, 2012; Iaria, Lanyon, Fox, Giaschi, & Barton, 2008; Maguire et al., 2000; Woollett & Maguire, 2011). However, in many of these studies, path integration, which contributes to topographical memory (Gallistel, 1990; Wang, 2016), might also have contributed to task performance, because landmarks were not isolated from path integration. Chrastil and colleagues recently reported that gray matter volume in the retrosplenial cortex, HPC, and medial prefrontal cortex positively predicted individual differences in path integration abilities (Chrastil, Sherrill, Aselcioglu, Hasselmo, & Stern, 2017). However, no studies have compared environmental-cue-navigation and path integration in the same spatial context and on the same participant sample. Such direct comparisons are crucial for controlling confounds like task demand, attentional engagement, individual idiosyncrasies, etc. Therefore, the anatomical correlates of path integration and environmental-cue-navigation – when the two processes are operating in the same spatial context – are not well understood at present.

HPC and entorhinal cortex (ERC) are essential to spatial navigation (Moser, Kropff, & Moser, 2008). These two brain structures seem to participate in both path integration and environmental-cue-navigation. For path integration, human fMRI studies have shown that HPC is recruited in this process (Chrastil, Sherrill, Hasselmo, & Stern, 2015; Wolbers, Wiener, Mallot, & Büchel, 2007), and that the strength of grid-cell-like activity in ERC is correlated with path integration ability in the elderly (Stangl et al., 2018). Consistently, animal studies have shown that self-motion cues influence both hippocampal place cells (Gothard, Skaggs, & McNaughton, 1996; Quirk, Muller, & Kubie, 1990) and entorhinal grid cells (Hafting, Fyhn, Molden, Moser, & Moser, 2005), and that lesions to HPC and ERC impair path integration performance (Parron & Save, 2004; Whishaw, McKenna, & Maaswinkel, 1997).

As to environmental-cue-navigation, evidence is relatively clear on the involvement of HPC and ERC when distal landmarks or geometric cues are concerned. Distal cues are usually positioned at a distance from the navigation space and are useful for providing directional information but not distance information (Lew, 2011). Animal studies have shown that hippocampal place cells and entorhinal grid cells are influenced by distal landmarks, in that their firing patterns closely followed the rotations of distal landmarks (Cressant, Muller, & Poucet, 1997; Hafting, Fyhn, Molden, Moser, & Moser, 2005; Muller & Kubie, 1987). Geometric cues refer to geometric boundaries, e.g., a square enclosure, or object arrays forming implicit geometric shapes (Cheng & Newcombe, 2005). Human MRI studies have shown that HPC is involved in topographical memory of complex landmark layouts (Hartley & Harlow, 2012; Maguire et al., 2000; Wolbers & Büchel, 2005), and that grid-cell-like activity in ERC is aligned with geometric boundary in visual space (Julian, Keinath, Frazzetta, & Epstein, 2018). Furthermore, rodent studies have shown that hippocampal place cells are controlled by enclosure shapes (Keinath, Julian, Epstein, & Muzzio, 2017), and that firing patterns of entorhinal grid cells are oriented to geometric boundaries in navigation space (Krupic, Bauza, Burton, Barry, & O’Keefe, 2015).

In contrast, evidence is controversial as to the neural basis of navigating with isolated local landmarks. In contrast to distal landmarks, local landmarks are usually positioned in or near the navigation space and can provide information on both direction and distance (Lew, 2011). When considered in isolation, a local landmark does not form any configural shape; hence, local landmarks are considered as featural cues, as opposed to geometric cues (Cheng & Newcombe, 2005). While some studies found that the processing of local landmarks, especially a single local landmark, was associative and was not related to hippocampal activity, compared to geometric and configural cues (Doeller, King, & Burgess, 2008; McDonald & White, 1994; Packard & McGaugh, 1992; Wegman, Tyborowska, & Janzen, 2014), rodent studies showed that hippocampal place cells respond to manipulations of an isolated landmark (Bjerknes, Dagslott, Moser, & Moser, 2018; Gothard et al., 1996) and display spatial modulations relative to local landmarks (Deshmukh & Knierim, 2013). These mixed findings suggest that navigation using local landmarks may depend on HPC. Moreover, rodent studies found that neurons in the lateral ERC displayed spatial specificity to local landmarks (Deshmukh & Knierim, 2011), suggesting that navigation using local landmarks might also recruit ERC.

Given the existing evidence for the involvement of HPC and ERC in environmental-cue-navigation and path integration, we hypothesized that anatomical variability, in particular gray matter volume, of these structures might be correlated with individual abilities to navigating with environmental cues and self-motion cues. To address these important questions, we used an individual differences approach, which allows for in-depth investigation of brain-behavior relationships by exploiting intersubject variability (Seghier & Price, 2018). In two experiments, we dissociated environmental-cue-navigation and path integration in the same navigation task and correlated performance with morphological variations. Given the existing ambiguous evidence on the involvement of HPC and ERC in local-landmark-navigation, we hoped to shed more light on this question by using a single isolated object as the local landmark in the navigation task.

## 2 EXPERIMENT 1

### 2.1 Methods

#### 2.1.1 Participants

The behavioral data of this experiment have been reported in a previous paper (Chen et al., 2017). Therefore, the method is only briefly described here. Twenty-four young healthy adults (12 female) from the local community of Magdeburg participated in the experiment. They ranged in age from 20 to 34 years old, with a mean of 25. All participants were right-handed, had normal or corrected-to-normal vision, and had no history of neurological diseases. Participants completed a navigation task in an immersive virtual reality environment on two consecutive days and underwent MRI structural scanning afterward on the third day. Participants also completed a battery of cognitive tests, the results of which were reported in Chen et al. (2017). Two participants were dropped from the analysis because they did not return for MRI scanning. This resulted in 22 participants (11 female, mean age = 25) in the analysis. All participants gave written informed consent and received monetary compensation. The experiment was approved by the local ethics committee of Otto-von-Guericke University, Magdeburg, Germany.

#### 2.1.2 Stimuli and task

The virtual environment was displayed via a Head Mounted Display (HMD, Oculus Rift Development Kit 2, Oculus VR LLC, http://www.oculus.com). Graphics were rendered using Vizard software (WorldViz, version 5, Santa Barbara, CA). The participant’s position was tracked using a Vicon motion tracking system (https://www.vicon.com) and the participant’s orientation was tracked via the inertial sensor of the HMD. Figure 1a and Figure 1b depict the virtual environment. The flag served as the local landmark. The navigation task was to walk through a series of three posts (outbound path) and attempt to walk back to the remembered location of the first post (inbound path) (Figure 1b). After participants had reached the third post, the environment disappeared, and participants counted backward from a number ranging from 100 to 200 in steps of 3 for 20 seconds. Next, participants were asked to walk back to the location of the first post. Finally, participants rated their confidence on navigation accuracy on a scale from 1 (very unconfident) to 10 (very confident), with intervals of 1.

**Figure 1:**
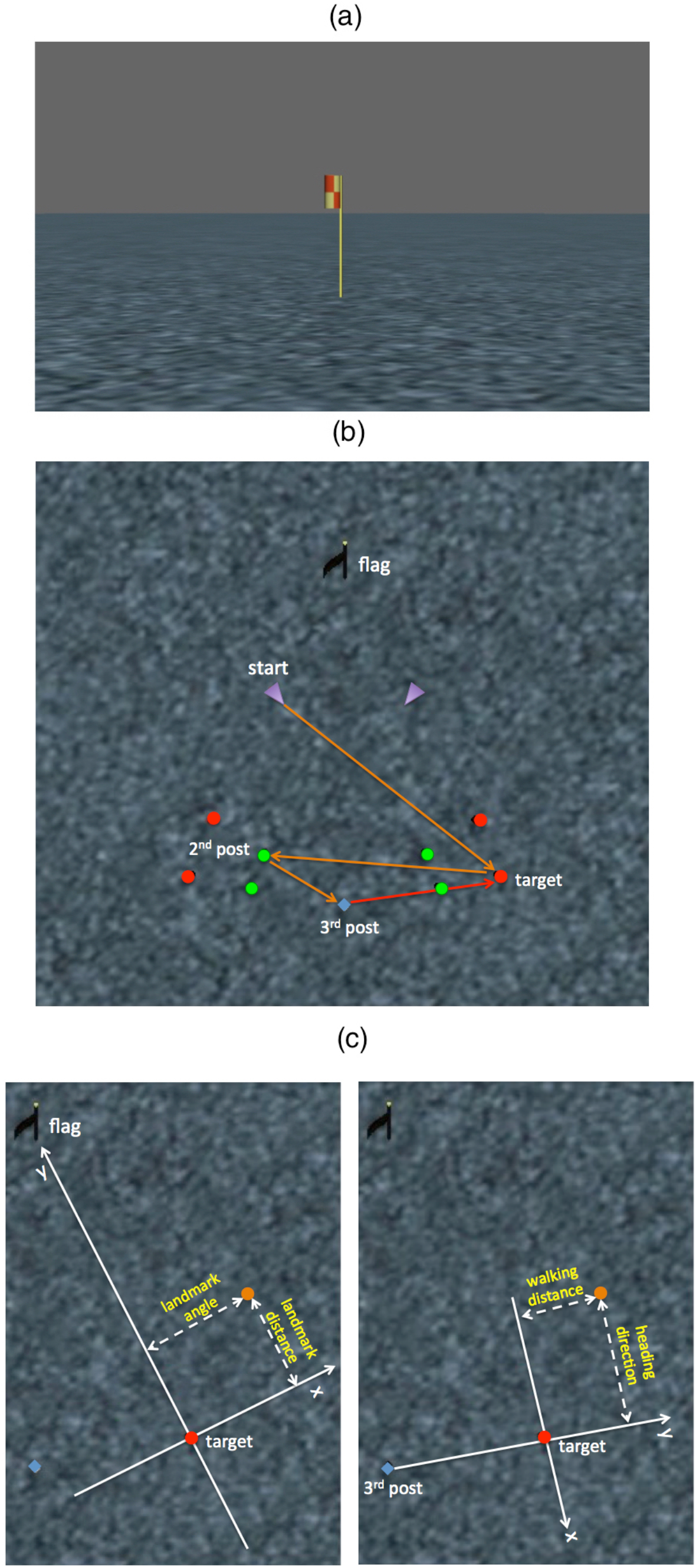
Experimental setup and behavioral analysis in Experiment 1. (a) a typical snapshot from the participant’s first-person perspective of the virtual environment. (b) a bird’s eye view of the environment. Purple triangles represent starting locations; red dots represent possible locations of the first post (i.e., target); green dots represent possible locations of the 2nd post; the blue diamond represents the location of the 3rd post. The solid orange and red arrows represent a typical trajectory of a trial: the orange arrows form the outbound path and the red arrow represents the correct inbound path. If we imagine a midline from the blue diamond to the flag, the first post always appeared on the opposite side to the location of the starting location in each trial. Randomized jitters, ranging from −0.4 m to 0.4 m in both spatial coordinates, were added to the first post’ location on a trial-by-trial basis (see Chen et al. 2017 for details). While standing at the starting location, the participant’s attention was directed to the flag by the experimenter before the outbound path started. During walking, participants were free to look at the flag. (c), spatial coordinates used to decompose the 2d distance error. The yellow dot represents the response location. In the landmark condition (left panel), the 2d distance error is decomposed into landmark-distance error and landmark-angle error in a spatial coordinate with the target location as the origin and the y-axis defined from the target location to the landmark location. In the self-motion condition (right panel), the 2d distance error is decomposed into heading-direction error and walking-distance error in a spatial coordinate with the target location as the origin and the y-axis defined from the 3^rd^ post to the target location.

The task could be completed with the landmark alone (landmark condition), self-motion cues alone (self-motion condition), with both cue types in congruence (combination condition), or with both cue types in incongruence (conflict condition). Importantly, the four conditions were identical during the outbound path – with the environment visible –, but they differed during the inbound path. Specifically, in the landmark condition, to eliminate the self-motion cues, participants were spun while sitting in a chair during the backward counting period. The visual world became visible during the inbound path, so that participants could rely on the landmark for localization. In the self-motion condition, participants stood still during counting, but the visual world and the landmark became invisible in the inbound path. Participants had to walk back in total darkness, relying on self-motion cues for localization. In the combination condition, participants remained oriented during counting, and the visual world became visible during the inbound path, so both landmark cues and self-motion cues were available for response. The conflict condition was similar to the combination condition, except that the landmark was located in a new position, which was 15° clockwise to its original location around the third post’s location. Since we were interested in contrasting landmark processing and self-motion processing, the analyses reported here focused on navigation performance in the two single-cue conditions. The behavioral task was administered on two consecutive days. On each day, there were 40 trials in total, each condition with 10 trials. Participants completed 4 practice trials at the beginning of each session. The task lasted about 1.5 hours each day.

#### 2.1.3 Behavioral analysis

In both the landmark condition and the self-motion condition, responses from different target posts were transformed into a common spatial coordinate. In the landmark condition, because participants relied on the landmark for response, the common spatial coordinate was defined in relation to the landmark location, with the target post’s location as the origin (0,0) and the direction from the target location to the landmark location as the y-axis (Figure 1c, left panel). Given previous studies suggesting differential mechanisms for distance and angular estimation (Allen, Kirasic, Rashotte, & Haun, 2004; Berthoz et al., 1999; Chrastil et al., 2017), we decomposed the 2-dimensional (2d) distance error into landmark-angle error along the x-axis and landmark-distance error along the y-axis (Figure 1c, left panel). This is consistent with the idea of landmark-vector representation, with the distance and angle components defined relative to the landmark (Collett, Cartwright, & Smith, 1986; Deshmukh & Knierim, 2013; McNaughton, Knierim, & Wilson, 1995). Although landmark-angle error was not directly measured as angles, it indicates how accurately participants retrieved the target’s orientation relative to the landmark. Landmark-distance error indicates how accurately participants retrieved the target’s distance to the landmark. Landmark-angle error and landmark-distance error were then correlated with brain morphology.

In the self-motion condition, because the landmark was invisible, and participants performed path integration, which started at the 3^rd^ post’s location, the common spatial coordinate was defined in relation to the 3^rd^ post’s location, with the target location as the origin and with the correct walking direction from the 3^rd^ post to the target location as the y-axis (Figure 1c, right panel). Analogous to the landmark condition, we decomposed the 2d distance error into heading-direction error along the x-axis and walking-distance error along the y-axis (Figure 1c, right panel). Although heading-direction error along the x-axis was not directly measured as angles, it is informative about how accurately participants retrieved the correct walking direction to the target location. Walking-distance error indicates how accurately participants retrieved the correct distance to walk to reach the target location. Heading-direction error and walking-distance error were then correlated with brain morphology. Therefore, in both coordinates in Figure 1c, x-axis and y-axis correspond to the angular component and distance component of the response error.

In both the landmark condition and the self-motion cue condition, trials from the two days were pooled together. To exclude outlier trials for each participant, in each condition, responses were excluded from the analysis if the 2d distance error were higher than the 3^rd^ quartile by more than 3 interquartile ranges.

#### 2.1.4 Image acquisition and pre-processing

After the participant had completed the navigation task, MR images were acquired in a 3T Siemens Prisma scanner on a separate day. A high-resolution whole-brain T1-weighted structural scan was acquired with the following MP-RAGE sequence: TR = 2500 ms, TE = 2.82 ms, flip angle = 7°, slices =192, orientation = sagittal, resolution = 1 mm isotropic. Since HPC and ERC can be clearly identified on T2-weighted images, a partial-volume high-resolution T2-weighted structural scan was acquired with the following MP-RAGE sequence: TR = 7930 ms, TE = 44 ms, flip angle = 180°, slices = 56, slice thickness = 1.1 mm, resolution = 0.4* 0.4*1.1 mm. The slices were acquired perpendicular to the long axis of HPC. If the T2-weighted structural scan did not have sufficient quality due to head motion, it was acquired for the second time. The scanning session took about 20-30 min. The T1-weighted image was bias-corrected in SPM12.

#### 2.1.5 Volumetry

To identify potential relationships between variability in brain morphology and navigational performance, we conducted volumetry analyses. In this analysis, regions of interest (ROIs), HPC and ERC, were manually segmented on coronal planes of the T2-weighted structural scan in ITK-SNAP (Version 3.4; www.itksnap.org; Yushkevich et al., 2006), using the segmentation protocol developed by Berron, Vieweg and colleagues (Berron et al., 2017). The full HPC and ERC were segmented from anterior to posterior. Anteriorly, the segmentation of ERC started four slices prior to the appearance of the hippocampal head; posteriorly, it ended 2 slices after the uncus had disappeared. The number of voxels of HPC and ERC was counted in ITK-SNAP, and their volumes were calculated as the number of voxels in the segmented region multiplied by the spatial resolution of the T2-weighted structural scan. The manual segmentation was performed by author XC, which was then checked and corrected by author PV. Both authors were blind to participants’ navigation performance. An example of manual segmentation is shown in Figure 2.

**Figure 2:**
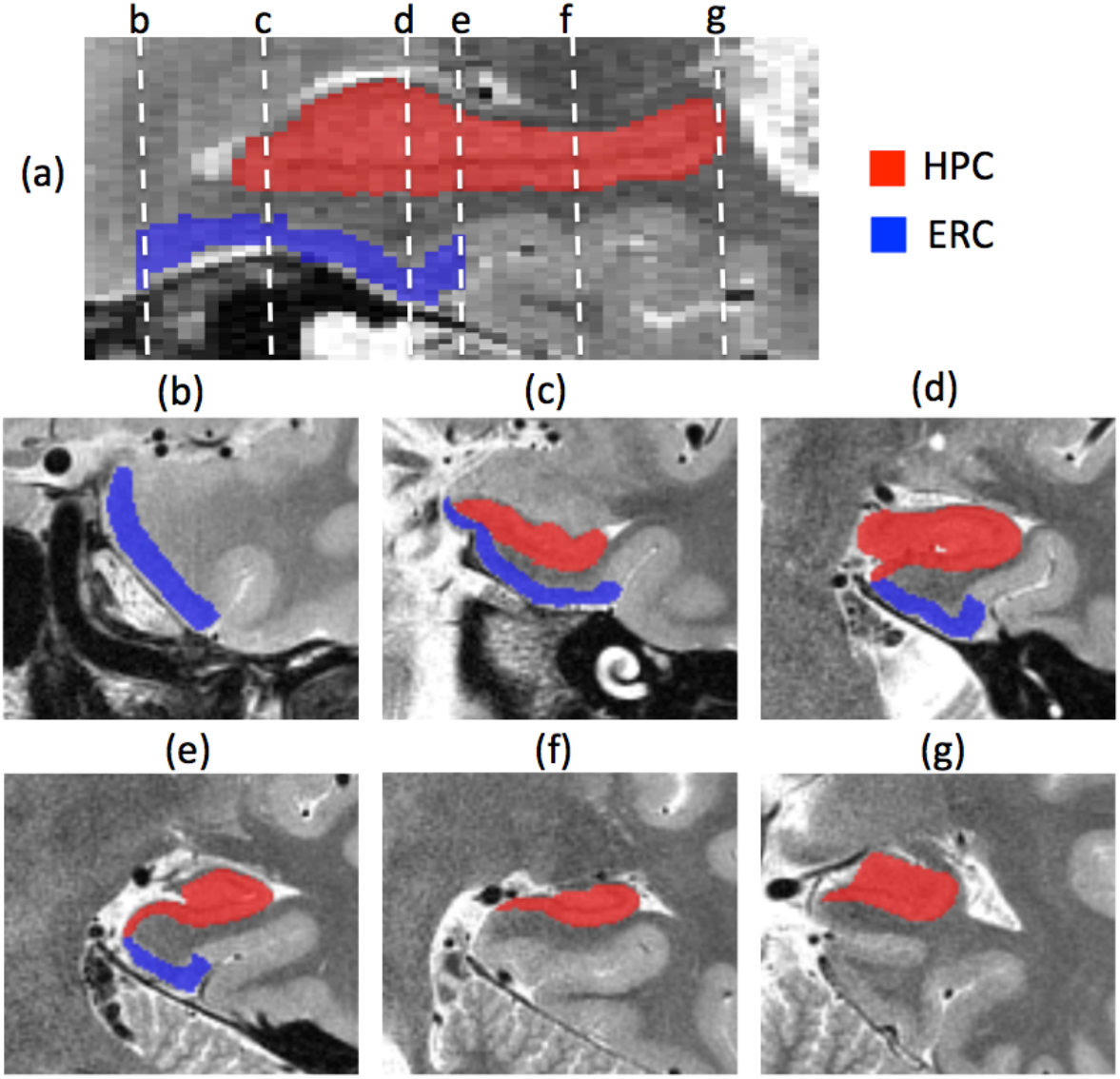
Demarcation of regions of interest. Manual segmentation of the hippocampus (HPC) and the entorhinal cortex (ERC) in the right hemisphere of a typical participant. (a), sagittal view of the brain. (b) to (g), cross-sections on the coronal plane corresponding to lines b to g in (a).

We then calculated total intra-cranial volume. In SPM12, we segmented the T1-weighted structural image into probabilistic tissue class images of gray matter, white matter, and cerebrospinal fluid (CSF). Total intra-cranial volume was equal to the sum of gray matter, white matter, and CSF. The ‘modulated’ non-linearly warped images were used (Malone et al., 2015). Total intra-cranial volume was used as a proxy of head size in the data analysis.

Finally, to determine whether anatomical variability was related to behavioral performance, we performed multivariate linear regression analyses, separately for the landmark condition and the self-motion condition, and separately for different dimensions of response errors. Because previous studies have shown hemispheric specificity of hippocampal and entorhinal functions (Kennepohl, Sziklas, Garver, Wagner, & Jones-Gotman, 2007; Maguire, Woollett, & Spiers, 2006; Schinazi, Nardi, Newcombe, Shipley, & Epstein, 2013; Shipton et al., 2014), the two hemispheres were analyzed separately. Covariates included in the regression analysis were HPC volume, ERC volume, head size (i.e., total intra-cranial volume), age, and sex, because we were interested in whether HPC volume or ERC volume contributed to individual differences in navigation performance, while controlling for head size, age, and sex. Response errors along different dimensions were the dependent variables.

### 2.2 Results

#### 2.2.1 Behavioral results

Response errors were analyzed in a 2 x 2 repeated-measures analysis of variance (ANOVA), with cue type (landmark vs. self-motion) and axis (angular vs. distance) as independent factors. Results are depicted in Figure 3. The main effect of cue type was not significant (F(1,21) = 0.395, p = 0.536, ŋ^2^ = 0.018), meaning that overall response error did not differ between cue types. The main effect of axis was significant (F(1,21) = 4.697, p = 0.042, ŋ^2^ = 0.183), meaning that overall participants committed larger angular errors than distance errors. The interaction between cue type and axis was significant (F(1,21) = 20.295, p < 0.001, ŋ^2^ = 0.491). Follow-up paired t tests showed that participants tended to commit larger landmark-distance error than landmark-angle error in the landmark condition, but the difference was not significant (t(21) = 1.675, p = 0.109, Cohen’s d = 0.357); participants committed larger heading-direction error than walking-distance error in the self-motion condition (t(21) = 5.836, p < 0.001, Cohen’s d = 1.244). Response errors in the self-motion condition were not correlated with response errors in the landmark condition (|rs| < 0.200, ps > 0.400), indicating a relative independence of landmark navigation and path integration. Women and men did not differ in any of the behavioral measurements (|ts| < 2.000, ps > 0.050). Age was not correlated with any of the behavioral measurements (|rs| < 0.100, ps > 0.750).

**Figure 3:**
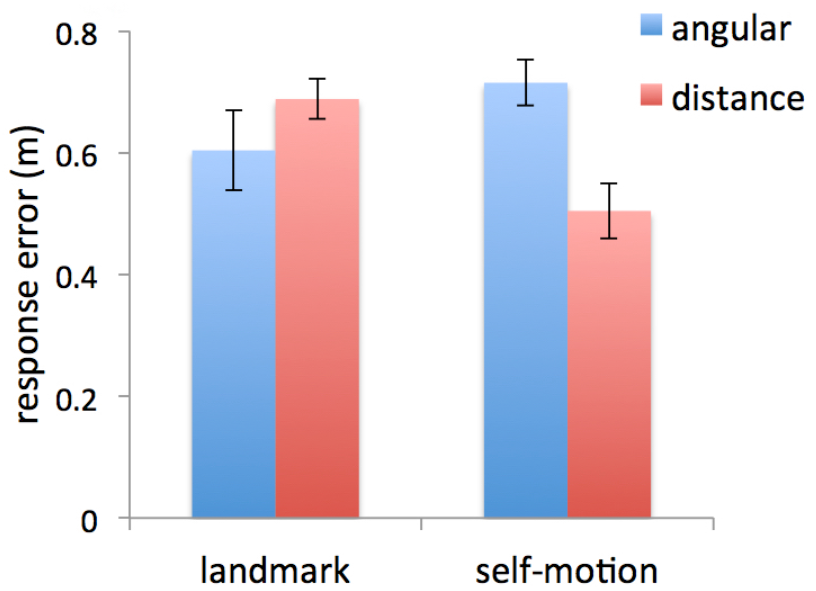
Behavioral results of Experiment 1. (a), response error is plotted as a function of cue type (landmark vs. self-motion) and axis (angular vs. distance). Angular error and distance error correspond to landmark-angle error and landmark-distance error respectively in the landmark condition, and correspond to heading-direction error and walking-distance error respectively in the self-motion condition. Error bars represent ±SE of the mean.

#### 2.2.2 MRI results

Table 1 lists bivariate correlations among predictors in multivariate linear regression models. To reiterate, analyses were conducted separately for each hemisphere, because previous studies have shown hemispheric specificity of hippocampal and entorhinal functions (Kennepohl et al., 2007; Maguire et al., 2006; Schinazi et al., 2013; Shipton et al., 2014). In each hemisphere, HPC volume and ERC volume were included as covariates of interest, and sex, age, head size were included as covariates of no interest in the multivariate linear regression model. Given that the correlation between ERC volume and HPC volume in the right hemisphere was very high (r = 0.809) and that HPC volume and ERC volume were of primary interest to us, for the right hemisphere, additional multivariate linear regression models were run with either HPC volume or ERC volume included (together with sex, age, and head size), to avoid a potential multicollinearity problem.

**Table 1:** Bivariate Pearson correlations among predictors in Experiment 1.

|  | sex | age | head size | left HPC | right HPC | left ERC |
| --- | --- | --- | --- | --- | --- | --- |
| age | -0.225 |  |  |  |  |  |
| head size | -0.657** | 0.015 |  |  |  |  |
| left HPC | -0.292 | -0.096 | 0.460* |  |  |  |
| right HPC | -0.288 | 0.093 | 0.583** | 0.734** |  |  |
| left ERC | 0.049 | 0.009 | 0.264 | 0.350 | 0.512* |  |
| right ERC | -0.384 | 0.323 | 0.464* | 0.520* | 0.809** | 0.435* |

\* represents $p < 0.05$ ; \*\* represents $p < 0.01$ .

##### 2.2.2.1 Predicting performance in the landmark condition

To reiterate, in the landmark condition, the 2d distance error was decomposed into landmark-distance error and landmark-angle error. Results are summarized in Table 2.

**Table 2:** Multivariate linear regression analyses for the landmark condition in Experiment 1.

| predictors | landmark-angle error |  |  | landmark-distance error |  |  |
| --- | --- | --- | --- | --- | --- | --- |
|  | beta | t | p | beta | t | p |
| left HPC | -0.169 | -0.596 | 0.560 | **-0.658** | **-2.753** | **0.014** |
| left ERC | 0.169 | 0.618 | 0.545 | 0.285 | 1.238 | 0.234 |
| sex | -0.005 | -0.015 | 0.988 | 0.05 | 0.170 | 0.867 |
| age | -0.124 | -0.490 | 0.631 | -0.107 | -0.502 | 0.623 |
| head size | 0.225 | 0.623 | 0.542 | 0.369 | 1.209 | 0.244 |
| | $F_{(5,16)} = 0.293, p = 0.910$ | | | $F_{(5,16)} = 1.716, p = 0.188$ | | |
| | $R^2 = 0.084, R^2_{adj} = -0.202$ | | | $R^2 = 0.349, R^2_{adj} = 0.141$ | | |
| right HPC | 0.032 | 0.068 | 0.947 | -0.501 | -1.064 | 0.303 |
|  | (-0.266) | (-0.913) | (0.374) | (-0.409) | (-0.439) | (0.168) |
| right ERC | -0.364 | -0.802 | 0.434 | 0.112 | 0.249 | 0.806 |
|  | (-0.340) | (-0.243) | (0.231) | (-0.264) | (-0.938) | (0.361) |
| sex | 0.056 | 0.164 | 0.872 | 0.298 | 0.88 | 0.392 |
| age | 0.019 | 0.073 | 0.943 | 0.022 | 0.083 | 0.935 |
| head size | 0.380 | 0.987 | 0.338 | 0.542 | 1.417 | 0.176 |
| | $F_{(5,16)} = 0.480, p = 0.786$ | | | $F_{(5,16)} = 0.524, p = 0.755$ | | |
| | $R^2 = 0.130, R^2_{adj} = -0.141$ | | | $R^2 = 0.141, R^2_{adj} = -0.128$ | | |

Multivariate linear regression analyses were run for landmark-angle error (left sections) and landmark-distance error (right sections) in the landmark condition separately, and for the left hemisphere (upper sections) and the right hemisphere (lower sections) separately. Significant regression coefficients are highlighted in red. Results when the right HPC and the right ERC were analyzed in separate regression models are shown in the parentheses.

For landmark-angle error, in the left hemisphere, when HPC volume and ERC volume were analyzed together, there were no significant correlations of either HPC volume (beta = −0.169, p = 0.560) or ERC volume (beta = 0.169, p = 0.545). The analysis on the right hemisphere revealed similar results when HPC volume and ERC volume were analyzed together (HPC, beta = 0.032, p = 0.947; ERC, beta = −0.364, p = 0.434). As mentioned previously, due to the high correlation between HPC volume and ERC volume in the right hemisphere, we did the same multivariate linear regression analyses for HPC and ERC in the right hemisphere separately, and the results remained unchanged (HPC, beta = −0.266, p = 0.374; ERC, beta = - 0.340, p = 0.231).

For landmark-distance error, in the left hemisphere, when HPC volume and ERC volume were analyzed together in the same multivariate linear regression model, HPC volume was negatively correlated with landmark-distance error (beta = −0.658, p = 0.014), meaning the larger the HPC, the smaller the error (Figure 4a); ERC volume was not correlated with landmark-distance error (beta = 0.285, p = 0.234). In the right hemisphere, when HPC volume and ERC volume were analyzed together, there were no significant correlations of HPC volume (beta = −0.501, p = 0.303) or ERC volume (beta = 0.112, p = 0.806). When HPC and ERC in the right hemisphere were analyzed separately, similar results were obtained (HPC, beta = −0.409, p = 0.168; ERC, beta = −0.264, p = 0.361).

**Figure 4:**
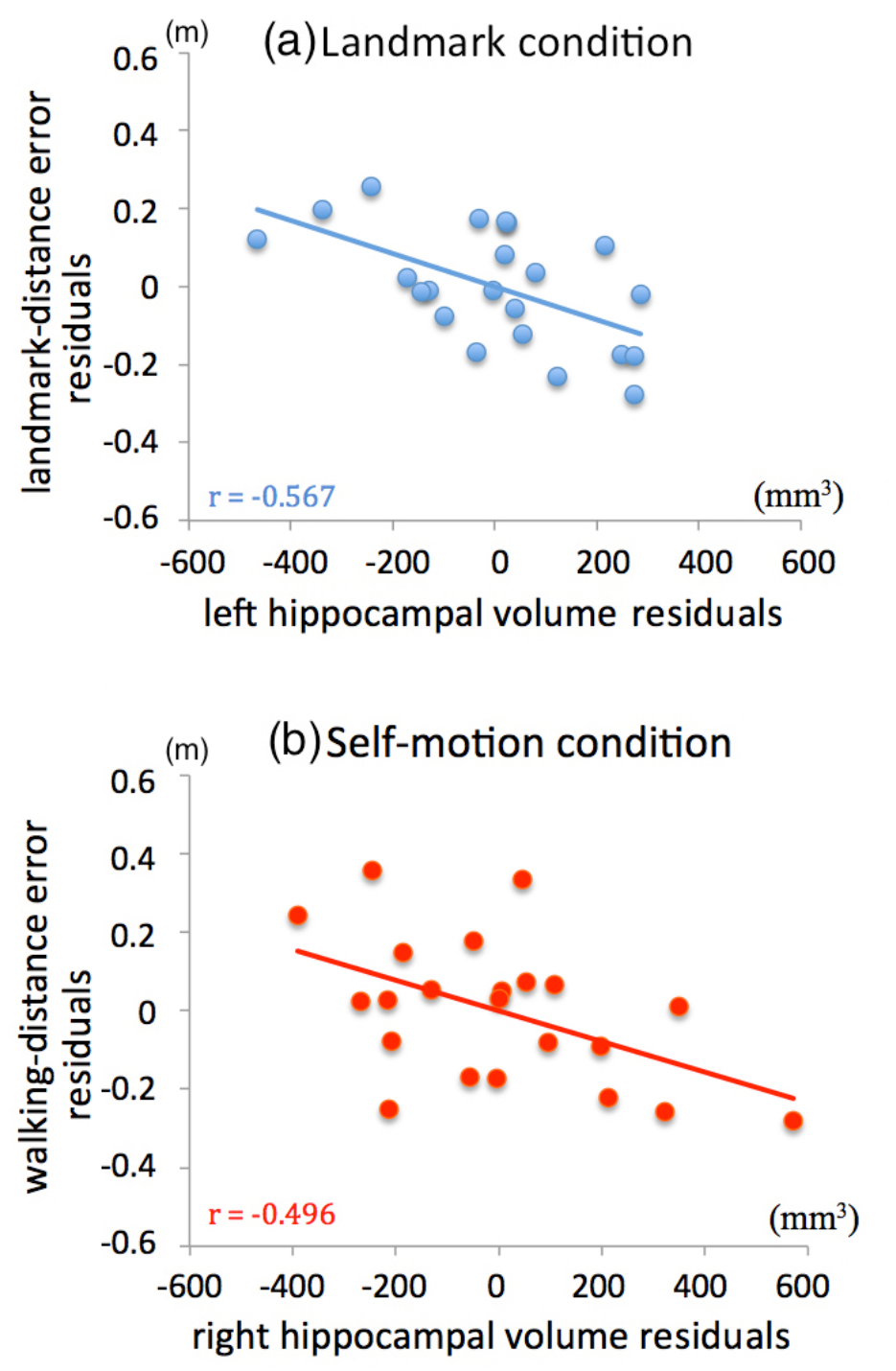
Partial regression plots in Experiment 1. The partial regression plot depicts the relationship between two variables after both of them are adjusted for other variables in the multiple regression model. For example, in (a), landmark-distance error and left HPC volume were regressed against all the other variables (i.e., age, sex, head size, and left ERC volume), from which residuals of the two dependent variables were obtained and plotted against each other. The partial regression plot in (b) was created in the same way. (a), the left HPC volume was correlated negatively with landmark-distance error in the landmark condition, after being adjusted for age, sex, head size, and left ERC volume. (b), the right HPC volume negatively predicted walking-distance error in the self-motion condition, after being adjusted for age, sex, and head size. The fitted linear regression line is displayed in each plot, together with the Pearson correlation coefficient (r) between the two adjusted variables.

To summarize, we found that in the landmark condition, HPC volume in the left hemisphere contributed positively to navigation performance. Specifically, participants with larger hippocampi performance better in terms of target-to-landmark distance. In contrast, ERC volume was not correlated with landmark navigation performance. We compared left HPC volume and left ERC volume in the analysis of landmark-distance error by looking at the 95% confidence intervals of beta weights (Cumming, 2009), and found beta weight differed significantly between the left HPC volume and the left ERC volume (p < 0.05). This suggests that the left HPC volume contributed to landmark navigation performance more substantially than the left ERC volume.

##### 2.2.2.2 Predicting performance in the self-motion condition

As described earlier, in the self-motion condition, the 2d distance error was decomposed into heading-direction error and walking-distance error. Results are shown in Table 3.

**Table 3:** Multivariate linear regression analyses for the self-motion condition in Experiment 1.

| predictors | heading-direction error |  |  | walking-distance error |  |  |
| --- | --- | --- | --- | --- | --- | --- |
|  | beta | t | p | beta | t | p |
| left HPC | -0.226 | -0.822 | 0.423 | -0.339 | -1.403 | 0.180 |
| left ERC | 0.003 | 0.01 | 0.992 | 0.026 | 0.112 | 0.912 |
| sex | 0.272 | 0.797 | 0.437 | 0.086 | 0.286 | 0.778 |
| age | 0.128 | 0.519 | 0.611 | -0.111 | -0.512 | 0.615 |
| head size | 0.063 | 0.179 | 0.860 | -0.276 | -0.895 | 0.384 |
| | $F_{(5,16)} = 0.504, p = 0.769$ | | | $F_{(5,16)} = 1.605, p = 0.215$ | | |
| | $R^2 = 0.136, R^2_{adj} = -0.134$ | | | $R^2 = 0.334, R^2_{adj} = 0.126$ | | |
| right HPC | -0.377 | -0.798 | 0.437 | -0.531 | -1.388 | 0.184 |
|  | (-0.097) | (-0.334) | (0.763) | (-0.542) | (-2.354) | (0.031) |
| right ERC | 0.343 | 0.757 | 0.460 | -0.014 | -0.038 | 0.970 |
|  | (0.060) | (0.214) | (0.833) | (-0.412) | (-1.766) | (0.095) |
| sex | 0.368 | 1.081 | 0.296 | 0.219 | 0.796 | 0.438 |
| age | 0.095 | 0.363 | 0.721 | 0.002 | 0.009 | 0.993 |
| head size | 0.084 | 0.217 | 0.831 | -0.023 | -0.075 | 0.941 |
| | $F_{(5,16)} = 0.493, p = 0.777$ | | | $F_{(5,16)} = 2.444, p = 0.079$ | | |
| | $R^2 = 0.133, R^2_{adj} = -0.137$ | | | $R^2 = 0.433, R^2_{adj} = 0.256$ | | |

Multivariate linear regression analyses were run for heading-direction error (left sections) and walking-distance error (right sections) in the self-motion condition separately, and for the left hemisphere (upper sections) and the right hemisphere (lower sections) separately. Significant regression coefficients are highlighted in red. Results when the right HPC and the right ERC were analyzed in separate regression models are shown in the parentheses.

For heading-direction error, analysis on the left hemisphere showed no significant correlations of HPC (beta = −0.226, p = 0.423) or ERC volume (beta = 0.003, p = 0.992) when the two were analyzed together in the same regression model. Similar results were obtained in the right hemisphere when HPC volume and ERC volume were analyzed together (HPC, beta = −0.377, p = 0.437; ERC, beta = 0.343, p = 0.460). The results remained the same when HPC and ERC in the right hemisphere were analyzed in separate regression models (HPC, beta = −0.097, p = 0.763; ERC, beta = 0.060, p = 0.833).

For walking-distance error, analysis on the left hemisphere revealed no correlations of HPC (beta = −0.339, p = 0.180) or ERC volume (beta = 0.026, p = 0.912) when the two were analyzed together in the same regression model. Similar results were obtained on the right hemisphere (HPC, beta = −0.531, p = 0.184; ERC, beta = −0.014, p = 0.970). However, when HPC and ERC in the right hemisphere were analyzed separately, HPC volume negatively correlated with walking-distance error (beta = −0.542, p = 0.031), meaning that the larger the HPC, the smaller the error (Figure 4b). Notably, this regression model significantly predicted walking-distance error (adjusted R^2^ = 0.300, F(4,17) = 3.245, p = 0.038). Regression coefficient of ERC volume remained non-significant when it was analyzed separately from HPC volume (beta = −0.412, p = 0.095). This suggests that the significant contribution of the right HPC volume to walking-distance error was obscured by its high correlation with the right ERC volume when both were included in the same regression model (variance inflation factors > 3.7).

To summarize, in the self-motion condition, HPC volume in the right hemisphere positively predicted navigation performance in terms of walking-distance to the target, meaning that participants with larger hippocampi retrieved the walking-distance to the target more correctly from a fixed location (the 3^rd^ post). Heading-direction error was not predicted by HPC volume. ERC volume did not predict any performance measurements in the self-motion condition.

### 2.3 Discussion

In Experiment 1, we found that hippocampal volume positively predicted navigation performance using both a local landmark and self-motion cues, but on different dimensions of responses. In the landmark condition, hippocampal volume was related to the accuracy of retrieving the target’s distance to the landmark but not the target’s orientation relative to the landmark. Different results on different dimensions could not be explained by statistical artifacts, such as range restriction, since the data were more dispersed along the dimension on which the correlation was absent (e.g., standard deviation was 0.153 m for landmark-distance error and 0.308 m for landmark-angle error). We also found that hippocampal volume was positively related to how accurately participants retrieved the walking distance to the target from a fixed location (i.e., the 3^rd^ post) in path integration trials. Again, statistical artifacts could not explain these results, since data dispersion was only slightly different between the two dimensions (standard deviation was 0.175 m for heading-direction error and 0.212 m for walking-distance error in the self-motion condition). Significant correlations between hippocampal volume and navigation performance were not observed in every dimension of behavioral measurements, indicating that our findings are unlikely to reflect general cognitive factors, such as memory capacity (Squire & Zola-Morgan, 1991) and attentional engagement. Rather, our findings suggest that hippocampal contribution to navigation performance is specific to the combination of cue type and response dimension.

The decomposition of the 2d distance error in the landmark condition into landmark-distance error and landmark-angle error rests on the assumption that participants localized the target location by extracting distance and angular information from visual features of the landmark, e.g., judging distance based on the visual size of the landmark (Gillam, 1995; Sedgwick, 1986). This corresponds to a view-based matching strategy (Collett et al., 1986). However, an alternative strategy available to participants was a reference orientation strategy: After the chair-spinning disorientation stage, participants could establish a reference orientation from self-position to the landmark, judge the orientation of the target location relative to the reference orientation, and then walk along the judged orientation. This strategy indicates that the response error in the landmark condition is better characterized by the spatial coordinate depicted in the Figure 1c, right panel, in which the 2d distance error is decomposed into heading-direction error and walking-distance error. Indeed, when applying this decomposition scheme to the landmark condition, we found that the left HPC negatively predicted heading-direction error (beta = −0.673, p = 0.004). This is not surprising given that geometrically, x-axis and y-axis are roughly switched between the two different coordinates in Figure 1c. This suggests that the correlation between HPC volume and landmark-distance error could actually be caused by the correlation between HPC volume and heading-direction error.

To analyze the possibility of the reference orientation strategy, we examined participants’ walking trajectories during response (i.e., the inbound path), when the participant’s instantaneous position was recorded every 0.1 seconds. We reasoned that if the reference orientation strategy was adopted throughout the response stage, the heading direction should be determined at the very early stage of the response, and the participant would then walk along the initially judged orientation until the response was made. Figure 5 shows a representative response trajectory. In this example, the participant’s heading direction was changed in the middle of the trajectory, indicating a change in strategy. The circuitousness of the 2^nd^ half of the trajectory indicates that the participant was finely adjusting self-position in an attempt to acquire a view of the landmark matched to the remembered visual snapshot taken during the outbound path. This implies that during response, if the participant adopted the reference orientation strategy, the participant did not rely on it exclusively. Instead, the participant also used the view-based matching strategy, especially in the later stage of response.

**Figure 5:**
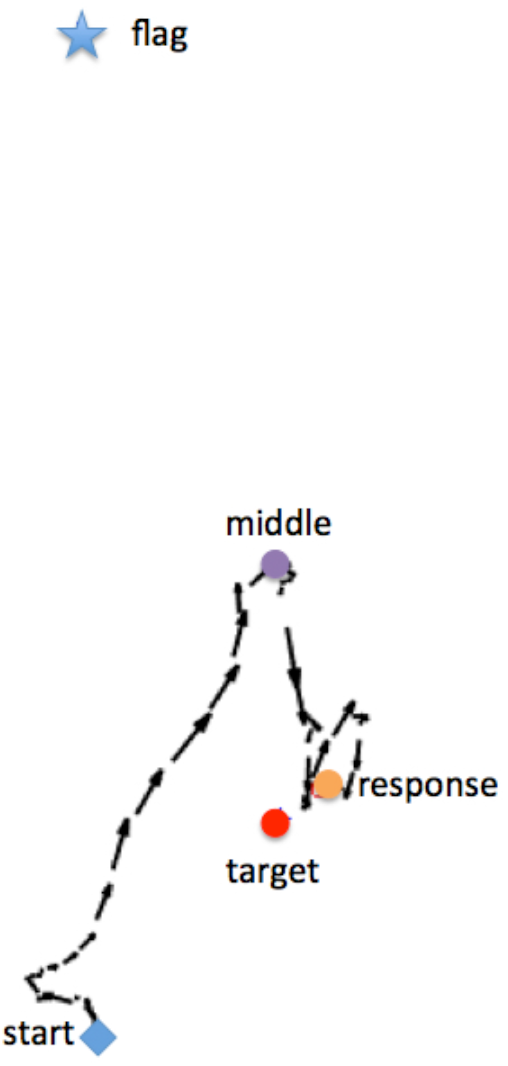
A representative response trajectory in Experiment 1. The black arrows represent the participant’s instantaneous positions, moving directions, and velocities. The arrow length represents speed. The blue diamond represents the starting position, the purple dot represents the middle position, the yellow dot represents the response location, the red dot represents the target location, and the blue star represents the landmark location. Note that the black arrows are not necessarily aligned with the facing direction of the participant.

To analyze the response trajectories quantitatively, for each trajectory, we obtained the middle position and the last position within the response time window and calculated their orientations relative to the 3^rd^ post. We found that the two orientations were only moderately correlated with each other across participants (r = 0.463, p = 0.030), suggesting that if participants initially used the landmark as a reference to specify the direction to walk along, they did not use this strategy throughout the entire response stage. We regressed heading-direction error at the middle position against HPC volume and ERC volume, with sex, age, and head size as covariates of no interest, and found that HPC volume in either hemisphere did not predict this behavioral measurement (ps > 0.150), suggesting that HPC volume was not related to how accurately people judged the target orientation relative to the reference orientation in the initial stage of response. We also added heading-direction error at the middle position as an additional predictor in the multivariate linear regression model for landmark-distance error; that is, landmark-distance error was regressed against heading-direction error at the middle position, in addition to HPC volume and ERC volume, with sex, age, and head size as covariates of no interest. The unique contribution of the left HPC volume to landmark-distance error remained significant (beta = −0.584, p = 0.022), suggesting that the correlation between HPC volume and landmark-distance error cannot be accounted for by the reference orientation strategy used in the initial stage of response. Finally, we simulated landmark-distance error assuming that participants had not changed the walking direction after the middle position and had walked the same distance from the 3^rd^ post to the response location. We did the same set of control analyses as we did for heading-direction error at the middle position. When the simulated landmark-distance error was regressed against HPC volume and ERC volume, with sex, age, and head size as covariates of no interest, HPC volume in either hemisphere did not predict the simulated landmark-distance error (ps > 0.290), indicating again that HPC volume was not related to how accurately people judged the target orientation relative to the reference orientation in the initial stage of response. When the simulated error was added as a predictor, in addition to HPC volume, ERC volume, sex, age, head size, into the multivariate linear regression model to predict landmark-distance error, the left HPC volume remained negatively correlated with landmark-distance error (beta = −0.551, p = 0.032), indicating again that the correlation between HPC volume and landmark-distance error cannot be accounted for by the reference orientation strategy used in the initial stage of response. Taken together, these results indicate that HPC volume was related to how accurately participants retrieved the target-to-landmark distance, which cannot be accounted for by the possible reference orientation strategy based on the landmark in the initial response stage.

Experiment 1 was conducted in an immersive virtual reality setup, in which full body-based cues were available. On the contrary, most previous studies looking at the anatomical or functional correlates of spatial navigation in humans were conducted in desktop virtual environments (e.g., Bohbot et al., 2007; Chrastil et al., 2017, 2015; Wolbers et al., 2007). We wondered whether the results of Experiment 1 could be replicated in a desktop virtual environment, in which body-based cues are absent and the sense of immersion is reduced. Therefore, in Experiment 2, we tried to conceptually replicate Experiment 1 in a desktop virtual reality setup. In addition, a different navigation task was used to further test the generalizability of the observations made in Experiment 1.

## 3 EXPERIMENT 2

While again dissociating the use of landmark vs. self-motion cues, we used a different navigation task in Experiment 2 and a desktop virtual reality setup. A single local landmark was used in a given trial in the landmark condition. In the self-motion condition, no landmarks were visible and optic flow served as self-motion cues for path integration, because physical translations and rotations were not possible. In both conditions, participants were required to memorize the location of a target, and they subsequently judged positions of test locations relative to the target location. Participants’ memory of the target location was tested on a linear track, with the local landmarks positioned on the linear track. Therefore, response error in the landmark condition reflected behavioral accuracy of retrieving the target’s distance to the landmark. This measurement was equivalent to landmark-distance error in the same condition in Experiment 1 (Figure 1c, left panel), in which significant correlation with hippocampal volume was observed. Response error in the self-motion condition was informative of behavioral accuracy of representing the target location in relation to the starting location where the movement started. This measurement was equivalent to walking-distance error in the self-motion condition in Experiment 1 (Figure 1c, right panel), in which significant correlation with hippocampal volume was observed. Therefore, Experiment 2 served as a conceptual replication of the effects observed in Experiment 1.

### 3.1 Method

#### 3.1.1 Participants

Twenty-four young healthy adults from the local community of Magdeburg participated in the experiment (15 male). These participants aged from 22 to 37 years old (mean = 27.9), were right-handed, had normal or corrected-to-normal vision, and had no history of neurological diseases. Two participants were excluded from the analysis because of low-quality T2-weighted structural scans caused by excessive head motion. This resulted in 22 participants in total in the data analysis (13 male, mean age = 27.9). All participants received monetary compensation and gave written informed consent to participate in the experiment. This experiment was approved by the local ethics committee of Otto-von-Guericke University, Magdeburg, Germany.

#### 3.1.2 Stimuli and task

Participants performed a relative location judgment task while lying inside the MRI scanner. Graphics were rendered using Vizard software (WorldViz, version 5, Santa Barbara, CA). The image was projected onto a screen mounted at the end of the MRI bed and reflected via an IR-reflecting first surface mirror. The display size was 22.9 cm in width and 12.9 cm in height. The distance from participants’ eyes to the screen was 100 cm.

Figure 6 shows the schematic of the environment layout. The task was to learn the location of a penguin, which was fixed in the environment. As shown in Figure 6a, two different trees were included, with one positioned closer to the penguin than the other. The distance from the close tree to the penguin was 9 m, and the distance from the distant tree to the penguin was 27 m. The close tree was 1.5 m tall, and the distant tree was 4.5 m tall. When standing at the penguin’s location, the vertical and horizontal visual angles subtended by the two trees were the same. There were three fixed starting points (position and orientation indicated by red arrows in Figure 6a). The distances from the penguin to each starting point was 8 m, and all the three arrows pointed to the penguin. In some trials, there was a collection of limited lifetime white dots (lifetime = 1 s) on the ground to provide optic flow information, as self-motion cues, during self-movement (Figure 6b, right & 6c, lower). In order to enhance the sense of spatial immersion, this layout was situated in a rich background environment, which was visible occasionally (Figure 6a, right). The background environment was an open-field area surrounded by mountains and trees. A lodge and a waterwheel were situated on the mountain.

**Figure 6:**
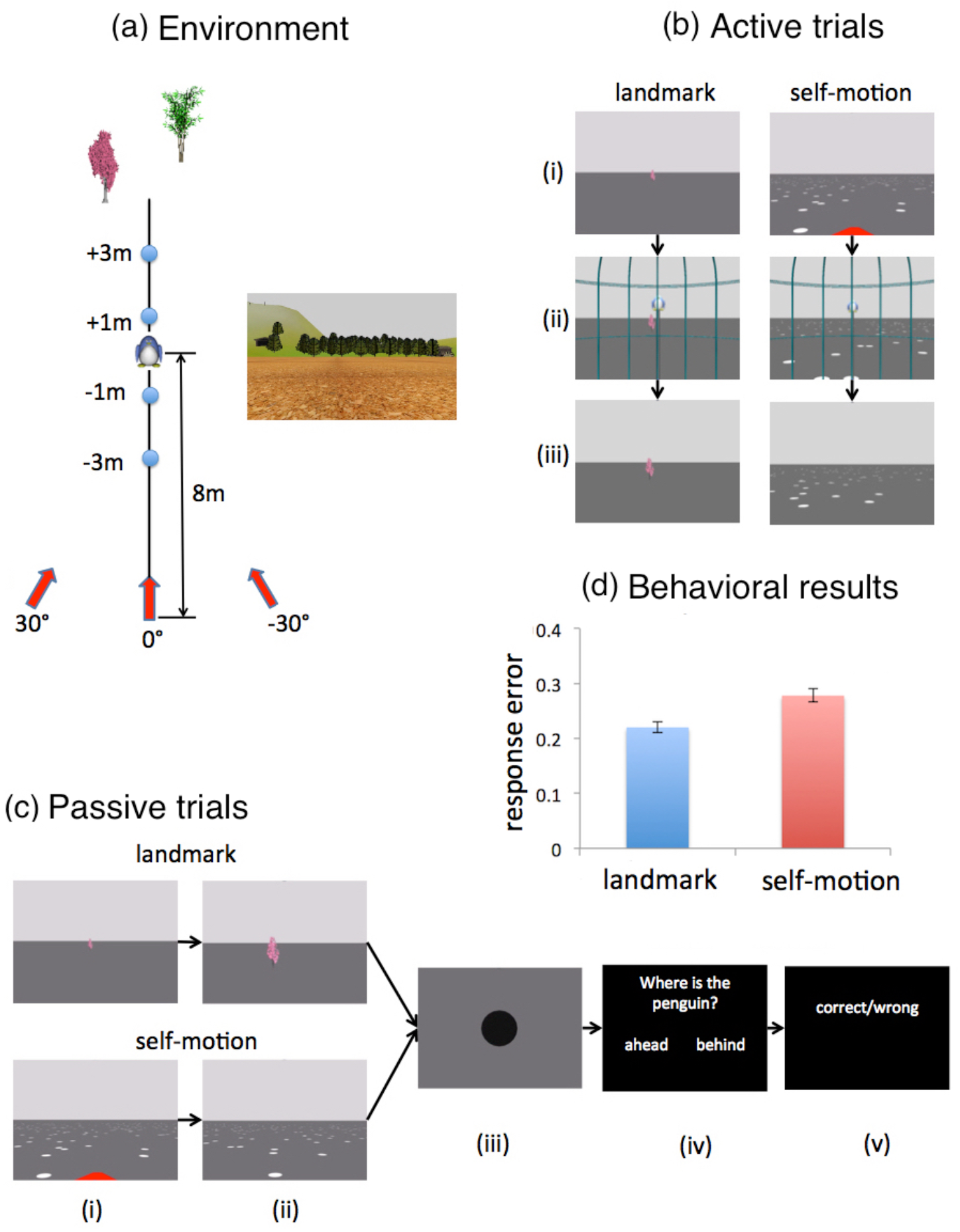
Experimental setup and behavioral results of Experiment 2. (a), left, schematic of the environment layout from a bird’s-eye view. For illustrative purposes, the landmarks are placed closer to the target than they really were. The target, which was a penguin, was 8 meters away from the three red arrows. The three red arrows pointed to the target along the 30°, 0°, and −30° perspectives, respectively. The four blue circles represent the four test locations — 3m behind (−3m), 1m behind (−1m), 1m ahead of (+1m), and 3m ahead of (+3m) the target, respectively. The four test locations lied on an imaginary linear line from the middle red arrow to the target. (a), right, a snapshot of the background environment. (b), the time course of active trials in the landmark condition (left) and the self-motion condition (right): (i) the participant stands at the start location, which was randomized in the landmark condition and fixed in the self-motion condition; (ii), the participant places the cage and the penguin appears at its correct location; (iii) the participant moves to the penguin’s location to make the penguin disappear. (c), the time course of passive trials in the landmark condition (upper) and the self-motion condition (lower): (i), the participant stands at the start location, which was randomized in the landmark condition and fixed in the self-motion condition; (ii), the participant is passively moved to the test location; (iii), at the test location, the subject’s viewpoint smoothly turns down and then faces vertically downward at the ground for 4 s; (iv), the participant judges the relative location of the penguin; (v), feedback is provided. The red arrow is invisible in the landmark condition and visible in the self-motion condition in stage (i) of both active and passive trials. (d), response error (= number of wrong trials / total number of trials) is plotted as a function of cue type. Error bars represent SE of the mean.

The task consisted of active trials and passive trials. In active trials (Figure 6b), participants learned the location of a target (a penguin, Figure 6a), and they ‘moved’ actively along a linear path by pressing appropriate buttons on a joystick. The participant first ‘moved’ to a location where they think the target was located, placing a cage at the location. Then, the target appeared at its correct location, providing feedback (Figure 6b). Finally, participants ‘moved’ to the correct location to correct the error. The movement could start from all three different starting points (red arrows in Figure 6a). The background environment was visible occasionally (Figure 6a). The sole purpose of active trials was to allow participants to learn the target location, and hence they were not analyzed further.

Passive trials tested participants’ memory of the target location. Passive trials represent the main part of the experiment and were analyzed. In passive trials (Figure 6c), participants were transported passively to a test location around the target, where the camera turned down and fixed at a large black dot on the blank ground for 4 s. Afterward, participants judged whether the penguin’s location was ahead or behind within 2 s. Feedback was provided afterward. The transportation always started from the 0° perspective, aligned with the pointing direction of the middle arrow (Figure 6a). Four different test locations were selected and were evenly spaced around the penguin’s location along the 0° perspective (Figure 6a). The penguin remained invisible throughout the trial. The background environment was never visible.

In both active trials and passive trials, the use of landmarks and the use of self-motion cues were dissociated. In the landmark condition (Figure 6b, left column & Figure 6c, upper row), the starting position of active movement in active trials and passive movement in passive trials was variable on a trial-by-trial basis. The starting position in each trial was randomly sampled from a uniform distribution [−4.5 m, 4.5 m] around the positions of the three fixed red arrows (Figure 6a). The movement was aligned with the pointing direction of the red arrow. The red arrows were invisible throughout the trial. In this way, participants would not have the knowledge of the distance they needed to travel to the target location and thus could not perform path integration to solve the task. In addition, the ground remained blank, and hence participants were forced to only rely on the landmark. The noise level of landmark cues was manipulated. Only the close tree was displayed in the landmark low-noise condition, and only the tree distant from the target was displayed in the landmark high-noise condition. Therefore, only one landmark was displayed in a given trial. Although the distant tree was farther away from the target location, it is still considered as a local landmark, since it was close enough to provide some degree of distance information, on which participants had to rely to complete the task. In passive trials, response error (= number of wrong trials / total number of trials; chance level = 0.5) was averaged across the two noise levels to get an estimate of how well participants utilized landmark cues for spatial localization overall.

In the self-motion condition, only self-motion cues were available (Figure 6b, right & Figure 6c, lower), while landmarks were invisible. The starting position of movement was fixed at one of the three red arrows in each trial, so participants knew in advance how far they needed to travel to reach the target. The movement direction was aligned with the pointing direction of the arrow, which was visible at the beginning of the trial. White limited lifetime dots (lifetime = 1 s) were displayed on the ground, providing optic flow information. The dots were moving along or against the moving direction of the participant. The noise level of self-motion cues was manipulated. In the self-motion low-noise condition, the movement speed of each dot was randomly sampled from a normal distribution with a mean of 0 m and a standard deviation of 0.2 m in the self-motion high condition. In the self-motion high-noise condition, the normal distribution had a mean of 0 m and a standard deviation of 6 m. Therefore, the white dots appeared to move considerably more in the self-motion high-noise condition than in the self-motion low-noise condition. As in the landmark conditions, in passive trials, response error was averaged across both noise levels to get an overall estimate of how well participant used self-motion cues for navigation. Distance judgment is considered a form of path integration (Chrastil et al., 2017; Jacob et al., 2017), since self-motion inputs need to be continuously integrated during the linear movement.

Finally, the experiment also comprised a compound condition, in which both cue types were available for self-localization. However, since our primary interest was to contrast landmark cues and path integration cues, this condition was not analyzed here.

#### 3.1.3 Procedure

First, participants were familiarized with the task and joystick operations outside of the scanner. While inside the scanner, participants first completed a practice block consisting of active trials and passive trials. Then they completed a test stage, which consisted of two runs. Each run had 5 blocks, corresponding to the compound condition, landmark low-noise condition, landmark high-noise condition, self-motion low-noise condition, and self-motion high-noise condition, randomized in order. In each block (except the compound condition block), participants first completed 5 active trials, followed by 20 passive trials. Each test location was visited 5 times in each block in passive trials. As mentioned above, only passive trials from landmark low-noise, landmark high-noise, self-motion low-noise, and self-motion high-noise conditions in the test stage were analyzed; response errors were averaged across the two noise levels as an index of overall navigational performance for landmark cues and self-motion cues separately.

#### 3.1.4 MRI data acquisition and preprocessing

Structural scans were acquired in a 7T MRI scanner (Siemens, Erlangen, Germany) at the Leibniz Institute for Neurobiology in Magdeburg with a 32-channel head coil (Nova Medical, Wilmington, MA). A high-resolution whole-brain T1-weighted structural scan was acquired with the following MP-RAGE sequence: TR = 1700 ms; TE = 2.01 ms; flip angle = 5°; slices = 176; orientation = sagittal; resolution = 1 mm isotropic. A partial-volume turbo spin echo high-resolution T2-weighted structural scan was acquired perpendicular to the long axis of the hippocampus with the following sequence: TR = 8000 ms; TE = 76 ms; flip angle = 60°; slices = 55; slice thickness = 1mm; distance factor = 10%; in-plane resolution = 0.4×0.4 mm; echo spacing = 15.1 ms, turbo factor = 9, echo trains per slice = 57. Structural scans were acquired while participants were performing the practice block. BOLD fMRI data were acquired while participants were performing the task in the test stage, but the results are not reported here. The T1-weighted structural scan was bias-corrected in SPM12.

#### 3.1.5 Volumetry

Following the same procedure as in Experiment 1 (section 2.1.5), HPC and ERC were manually segmented on T2-weighted structural scans, and their volumes were calculated and analyzed in multivariate linear regression models to predict navigation performance.

### 3.2 Results

#### 3.2.1 Behavioral results

Behavioral results are shown in Figure 6d. Similar to Experiment 1, response errors were not correlated between landmark cues and self-motion cues (r = 0.198, p = 0.376), indicating that navigation with these two different types of spatial cues might involve relatively independent mechanisms. Paired t-tests showed that participants performed better with landmark cues than self-motion cues (t(21) = 4.067, p = 0.001, Cohen’s d = 0.867). This was expected considering that human navigators are generally poor in path integration and that body-based cues, which are critical for path integration, were absent in the current experiment. Independent-sample t tests showed that women and men did not differ on navigation performance with landmark cues (t(20) = 1.949, p = 0.065, Cohen’s d = 0.845) or self-motion cues (t(20) = 0.117, p = 0.908, Cohen’s d = 0.051). Age was not correlated with navigation performance with landmark cues (r = 0.123, p = 0.586) or self-motion cues (r = −0.214, p = 0.338).

#### 3.2.2 MRI analyses

Table 4 lists bivariate correlations among predictors in multivariate linear regression models. Similar to Experiment 1, analyses were conducted separately for each hemisphere, with HPC volume and ERC volume as covariates of interest, and sex, age, head size as covariates of no interest.

**Table 4:** Bivariate Pearson correlations among predictors in Experiment 2.

|  | sex | age | head size | left HPC | right HPC | left ERC |
| --- | --- | --- | --- | --- | --- | --- |
| age | -0.108 |  |  |  |  |  |
| head size | -0.680** | 0.031 |  |  |  |  |
| left HPC | -0.476* | -0.115 | 0.771** |  |  |  |
| right HPC | -0.534* | -0.129 | 0.796** | 0.985** |  |  |
| left ERC | -0.234 | -0.139 | 0.334 | 0.426* | 0.416 |  |
| right ERC | -0.172 | -0.198 | 0.354 | 0.222 | 0.234 | 0.378 |

\* represents $p < 0.05$ ; \*\* represents $p < 0.01$ .

##### 3.2.2.1 Predicting performance in the landmark condition

We conducted multivariate linear regression analyses to predict response error in the landmark condition (i.e., landmark-distance error). Results are summarized in Table 5. The analysis on the left hemisphere showed that when HPC and ERC were analyzed together, HPC volume contributed negatively to landmark-distance error (beta = −0.997, p = 0.002). As shown in Figure 7a, participants with larger hippocampi committed smaller errors with the landmark cue. ERC volume did not predict landmark-distance error (beta = −0.019, p = 0.918). Notably, this regression model significantly predicted landmark-distance error as a whole (adjusted R^2^ = 0.467, F(5,16) = 4.687, p = 0.008).

**Table 5:**
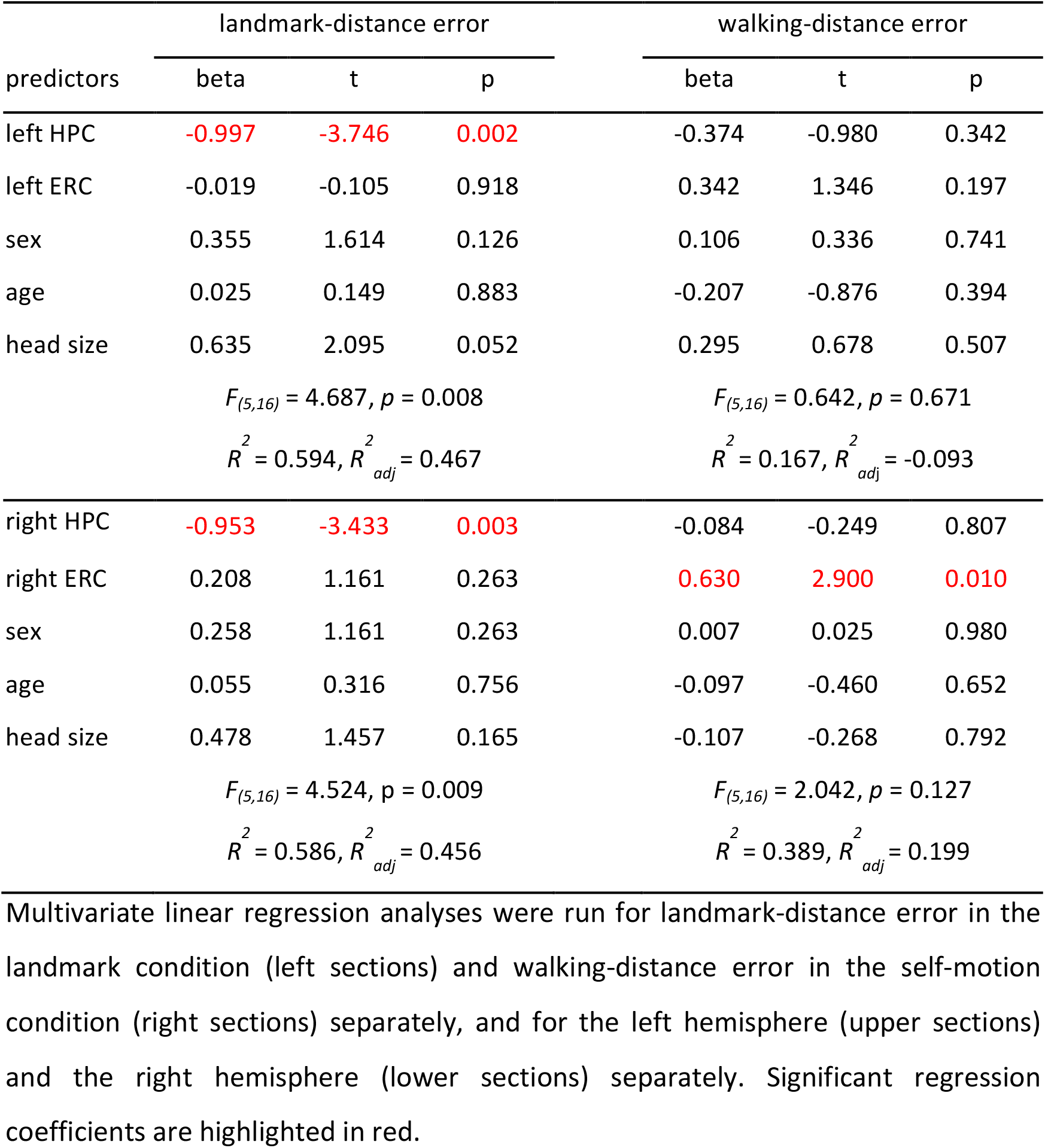
Multivariate linear regression analyses in Experiment 2.

| predictors | landmark-distance error |  |  | walking-distance error |  |  |
| --- | --- | --- | --- | --- | --- | --- |
|  | beta | t | p | beta | t | p |
| left HPC | **-0.997** | **-3.746** | **0.002** | -0.374 | -0.980 | 0.342 |
| left ERC | -0.019 | -0.105 | 0.918 | 0.342 | 1.346 | 0.197 |
| sex | 0.355 | 1.614 | 0.126 | 0.106 | 0.336 | 0.741 |
| age | 0.025 | 0.149 | 0.883 | -0.207 | -0.876 | 0.394 |
| head size | 0.635 | 2.095 | 0.052 | 0.295 | 0.678 | 0.507 |
| | $F_{(5,16)} = 4.687, p = 0.008$ | | | $F_{(5,16)} = 0.642, p = 0.671$ | | |
| | $R^2 = 0.594, R^2_{adj} = 0.467$ | | | $R^2 = 0.167, R^2_{adj} = -0.093$ | | |
| right HPC | **-0.953** | **-3.433** | **0.003** | -0.084 | -0.249 | 0.807 |
| right ERC | 0.208 | 1.161 | 0.263 | **0.630** | **2.900** | **0.010** |
| sex | 0.258 | 1.161 | 0.263 | 0.007 | 0.025 | 0.980 |
| age | 0.055 | 0.316 | 0.756 | -0.097 | -0.460 | 0.652 |
| head size | 0.478 | 1.457 | 0.165 | -0.107 | -0.268 | 0.792 |
| | $F_{(5,16)} = 4.524, p = 0.009$ | | | $F_{(5,16)} = 2.042, p = 0.127$ | | |
| | $R^2 = 0.586, R^2_{adj} = 0.456$ | | | $R^2 = 0.389, R^2_{adj} = 0.199$ | | |

Multivariate linear regression analyses were run for landmark-distance error in the landmark condition (left sections) and walking-distance error in the self-motion condition (right sections) separately, and for the left hemisphere (upper sections) and the right hemisphere (lower sections) separately. Significant regression coefficients are highlighted in red.

**Figure 7:**
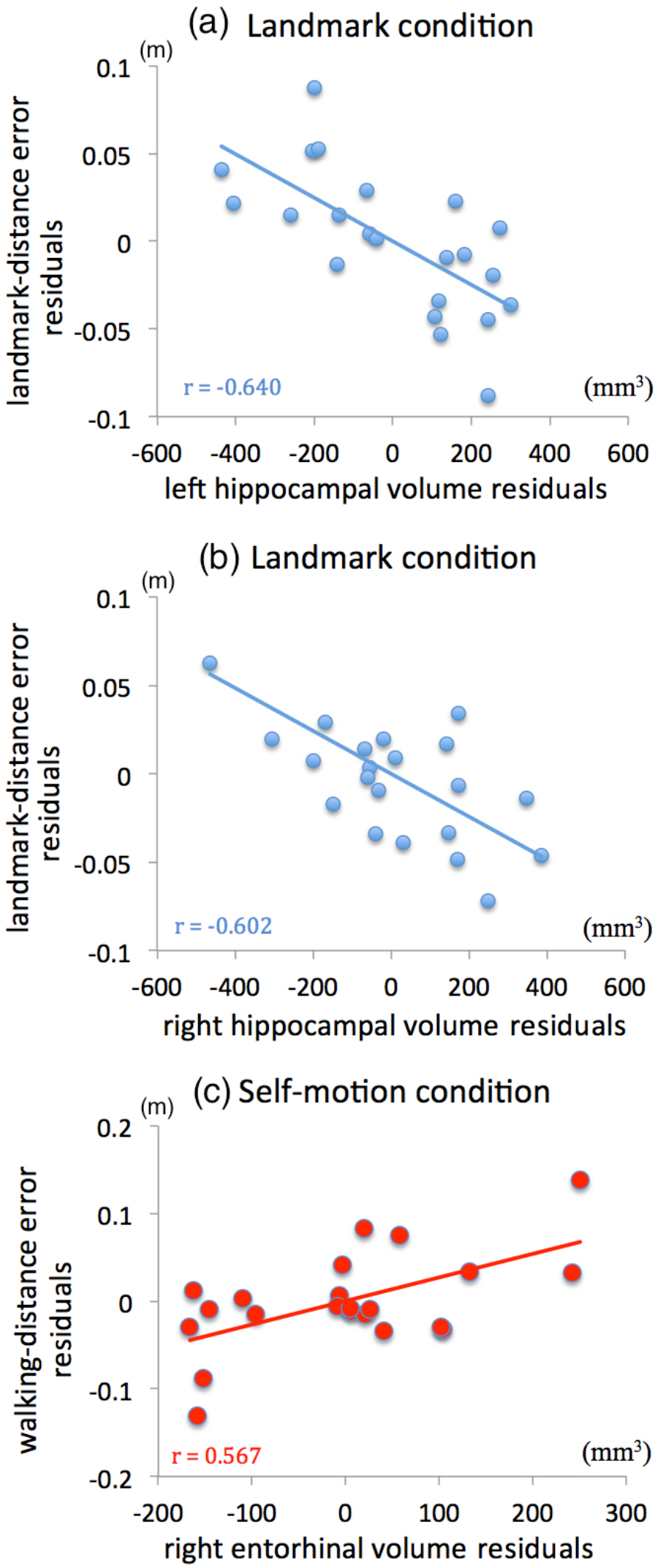
Partial regression plots in Experiment 2. As in Figure 4, variables in each plot were adjusted for other variables in the multivariate linear regression model. HPC volume negatively predicted landmark-distance error in the landmark condition in the left hemisphere (a) and the right hemisphere (b), after both variables were adjusted for age, sex, head size, and ERC volume in the corresponding hemisphere. (c), the right ERC volume positively predicted walking-distance error in the self-motion condition, after both variables were adjusted for age, sex, head size, and the right HPC volume. In each plot, the fitted linear regression line is displayed together with the Pearson correlation coefficient (r) between the two adjusted variables.

Similar results were obtained on the right hemisphere. When HPC and ERC were analyzed together, HPC volume contributed negatively to landmark-distance error (beta = −0.953, p = 0.003). As shown in Figure 7b, the larger the right HPC volume, the smaller the error. In contrast, ERC volume did not predict landmark-distance error (beta = 0.208, p = 0.263). Notably, this regression model significantly predicted landmark-distance error as a whole (adjusted R^2^ = 0.456, F(5,16) = 4.524, p = 0.009).

To summarize, the results showed that hippocampal volume in both hemispheres positively contributed to individual differences in navigation performance with a single local landmark. Specifically, participants with larger hippocampi retrieved more correctly the target’s distance to the local landmark. ERC volume did not contribute to landmark navigation performance. By comparing the 95% confidence intervals of beta weights (Cumming, 2009), we found that beta weight differed significantly between HPC volume and ERC volume in both hemispheres (ps < 0.05). This pattern of results is consistent with the findings in Experiment 1.

##### 3.2.2.2 Predicting performance in the self-motion condition

We conducted multivariate linear regression analyses to predict response error in the self-motion condition (i.e., walking-distance error). Results are summarized in Table 5.

Analysis on the left hemisphere revealed that neither HPC volume nor ERC volume was correlated with walking-distance error (HPC, beta = −0.374, p = 0.342; ERC, beta = 0.342, p = 0.197) when they were analyzed together.

Analysis on the right hemisphere revealed that when HPC and ERC were analyzed together, HPC volume did not predict walking-distance error (beta = −0.084, p = 0.807); ERC volume positively predicted walking-distance error (beta = 0.630, p = 0.010), meaning that participants with larger ERC performed worse with self-motion cues (Figure 7c).

To summarize, the results showed that HPC volume was not correlated with path integration performance, and ERC volume in the right hemisphere negatively predicted path integration performance. Specifically, participants with larger ERC performed worse in terms of walking distance to the target location.

### 3.3 Discussion

In Experiment 2, results in the landmark condition were very similar to those in the landmark condition in Experiment 1. Hippocampal volume positively predicted performance with the single local landmark, meaning the larger the hippocampus, the more accurate the retrieved distance from the target to the landmark. Unlike Experiment 1, we did not observe correlations between hippocampal volume and path integration performance, indicating that hippocampal involvement in path integration might be stronger when body-based cues were available. The absence of correlation between hippocampal volume and path integration performance also indicates that the significant hippocampal contribution to landmark navigation performance cannot be purely explained by general cognitive abilities, such as memory capacity (Squire & Zola-Morgan, 1991) or attentional engagement. Finally, the absence of hippocampal correlations with path integration performance could not be explained by statistical artefacts either, since we did observe a significant correlation between entorhinal volume and path integration performance.

## 4 POOLED ANALYSIS

As described previously, Experiment 1 and Experiment 2 shared two performance measurements, landmark-distance error in the landmark condition and walking-distance error in the self-motion condition. Therefore, to test the robustness of the observed effects, we pooled the data across the two experiments and conducted the same set of multivariate linear regression analyses on all the 44 participants. Before the data pooling, performance measurements and ROI volumes were standardized, so z scores were included in the regression analyses. Performance measurements were standardized because they were calculated from different navigation tasks and on different scales in the two experiments. ROI volumes were standardized because T2-weighted structural scans obtained in different MRI scanners had different contrasts (although the spatial resolution was the same), which could have affected manual segmentation. Age, sex, and head size were not standardized, since they were already on the same scale across experiments.

Table 6 lists bivariate correlations among predictors. For each hemisphere, HPC and ERC were analyzed together in the same regression models. Given that the correlation between HPC volume and ERC volume in the right hemisphere was again substantially high (r = 0.613), for the right hemisphere, we also analyzed HPC and ERC in separate regression models to avoid a potential multicollinearity problem.

**Table 6:** Bivariate correlations among predictors in the pooled analysis.

|  | sex | age | head size | Z(left HPC) | Z(right HPC) | Z(left ERC) |
| --- | --- | --- | --- | --- | --- | --- |
| age | -0.183 |  |  |  |  |  |
| head size | -0.649** | -0.022 |  |  |  |  |
| Z(left HPC) | -0.382* | -0.099 | 0.613** |  |  |  |
| Z(right HPC) | -0.408** | -0.025 | 0.685** | 0.859** |  |  |
| Z(left ERC) | -0.061 | -0.096 | 0.307* | 0.286 | 0.373* |  |
| Z(right ERC) | -0.308* | 0.067 | 0.395** | 0.473** | 0.613** | 0.406** |

ROI volumes were standardized within each experiment (z-score). \* represents $p < 0.05$ ; \*\* represents $p < 0.01$ .

### 4.1 ROI volumetry analyses

#### 4.1.1 Predicting landmark-distance error in the landmark condition

The results are summarized in Table 7. In the left hemisphere, when HPC and ERC were analyzed together, HPC volume negatively predicted landmark-distance error (beta = −0.767, p < 0.001), meaning that participants with larger hippocampi committed smaller errors (Figure 8a); ERC volume did not predict landmark-distance error (beta = 0.243, p = 0.073). Notably, the regression model significantly predicted landmark-distance error as a whole (adjusted R^2^ = 0.360, F(5,38) = 5.843, p < 0.001).

**Table 7:** Multivariate linear regression analyses in the pooled analysis.

| predictors | landmark-distance error |  |  | walking-distance error |  |  |
| --- | --- | --- | --- | --- | --- | --- |
|  | beta | t | p | beta | t | p |
| left HPC | **-0.767** | **-4.901** | **< 0.001** | -0.259 | -1.384 | 0.174 |
| left ERC | 0.243 | 1.846 | 0.073 | **0.323** | **2.048** | **0.048** |
| sex | 0.192 | 1.138 | 0.262 | < 0.001 | < 0.001 | > 0.999 |
| age | 0.033 | 0.258 | 0.798 | -0.150 | -0.987 | 0.330 |
| head size | 0.381 | 1.959 | 0.057 | -0.159 | -0.683 | 0.499 |
| | $F_{(5,38)} = 5.843, p < 0.001$ | | | $F_{(5,38)} = 1.792, p = 0.138$ | | |
| | $R^2 = 0.435, R^2_{adj} = 0.360$ | | | $R^2 = 0.191, R^2_{adj} = 0.084$ | | |
| right HPC | **-0.687** | **-3.083** | **0.004** | -0.418 | -1.725 | 0.093 |
|  | **(-0.637)** | **(-3.376)** | **(0.002)** | **(-0.326)** | **(-1.585)** | **(0.121)** |
| right ERC | 0.078 | 0.439 | 0.663 | 0.140 | 0.730 | 0.470 |
|  | **(-0.206)** | **(-1.233)** | **(0.225)** | **(-0.032)** | **(-0.190)** | **(0.850)** |
| sex | 0.297 | 1.566 | 0.126 | 0.109 | 0.530 | 0.599 |
| age | 0.085 | 0.588 | 0.560 | -0.150 | -0.962 | 0.342 |
| head size | **0.496** | **2.136** | **0.039** | 0.083 | 0.329 | 0.744 |
| | $F_{(5,38)} = 2.835, p = 0.029$ | | | $F_{(5,38)} = 1.256, p = 0.303$ | | |
| | $R^2 = 0.272, R^2_{adj} = 0.176$ | | | $R^2 = 0.075, R^2_{adj} = -0.020$ | | |

Multivariate linear regression analyses were run for landmark-distance error in the landmark condition (left sections) and walking-distance error in the self-motion condition (right sections) separately, and for the left hemisphere (upper sections) and the right hemisphere (lower sections) separately. Performance measurements and ROI volumes were standardized within each experiment (z-score). Significant regression coefficients are highlighted in red.

**Figure 8:**
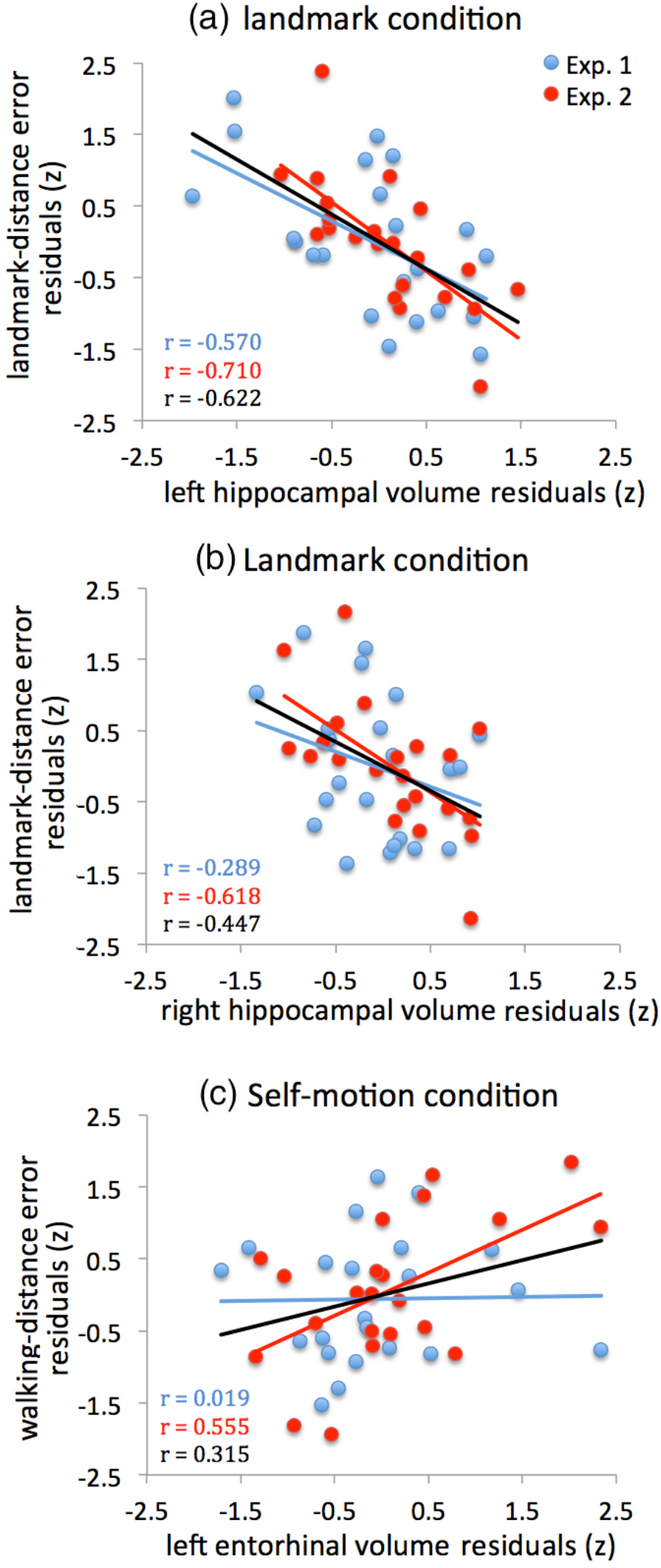
Partial regression plots in the ROI volumetry analysis on the pooled data. Performance measurements and ROI volumes were standardized (z-scores) in each experiment before pooling. As in Figure 4 and Figure 7, in each plot, variables were adjusted for other variables in the multivariate linear regression model. The left HPC volume (a) and the right HPC volume (b) were negatively correlated with landmark-distance error in the landmark condition. (c), the ERC volume positively predicted walking-distance error in the self-motion condition. Data points are displayed separately for each experiment (blue symbols for Experiment 1, red symbols for Experiment 2). Fitted linear regression lines are displayed for the two experiments separately (blue line and red line) and for the pooled data (dark line). Pearson correlation coefficients between adjusted variables (r) for individual experiments (blue and red texts) and pooled data (dark text) are indicated.

Similar results were obtained in the right hemisphere. When HPC and ERC were analyzed together, HPC volume negatively predicted landmark-distance error (beta = −0.687, p = 0.004), meaning the larger the HPC, the smaller the error (Figure 8b); ERC volume did not predict performance error (beta = 0.078, p = 0.663). Notably, this regression model significantly predicted landmark-distance error as a whole (adjusted R^2^ = 0.176, F(5,38) = 2.835, p = 0.029). Results remained unchanged when HPC and ERC in the right hemisphere were analyzed in separate regression models (HPC, beta = −0.637, p = 0.002; ERC, beta = −0.206, p = 0.225).

#### 4.1.2 Predicting walking-distance error in the self-motion condition

The results are summarized in Table 7. In the left hemisphere, when HPC and ERC were analyzed together, HPC volume did not predict walking-distance error (beta = −0.259, p = 0.174). ERC volume positively predicted walking-distance error (beta = 0.323, p = 0.048), meaning the larger the ERC, the larger the error (Figure 8c). Scatterplot in Figure 8c suggests that this negative correlation was mainly driven by data from Experiment 2, since the fitted linear regression line was almost flat in Experiment 1. In the right hemisphere, neither HPC volume nor ERC volume predicted walking-distance error (HPC, beta = −0.418, p = 0.093; ERC, beta = 0.140, p = 0.470) when they were analyzed together. Results remained unchanged when HPC and ERC in the right hemisphere were analyzed in separate regression models (HPC, beta = −0.326, p = 0.121; ERC, beta = −0.032, p = 0.850).

### 4.2 Voxel-based morphometry (VBM) analysis on pooled data

We conducted VBM analysis, which examines local morphological variability at the voxel level, to complement the ROI volumetry analyses. VBM can help verify results obtained in the volumetry analysis, because ROI demarcation in volumetry analysis may contain inaccuracies. Furthermore, VBM can reveal effects not detectable in the volumetry analysis, because an effect localized to a certain portion of a ROI could be washed out when the ROI is considered as a whole. Finally, unlike the volumetry analysis, VBM can examine the entire brain and capture effects beyond the ROI. In brief, in the VBM analysis, gray matter images of individual participants were registered to the Montreal Neurological Institute (MNI) template, and then a group-level general linear model was conducted to assess whether local gray matter volume at each voxel was correlated with individual differences in navigation performance.

The VBM analysis was conducted using the Dartel toolbox as implemented in SPM12 (Ashburner, 2007). First, T1-weighted structural scans were manually reoriented and shifted to approximately align with the MNI template. Second, T1-weighted structural scans were segmented into grey matter, white matter, and cerebrospinal fluid (CSF). Third, a population template was created as follows: an average brain of the group was created, and the deformations for individual images relative to the average brain were estimated and used to register individual images to the average brain; this procedure was conducted in an iterative manner. Fourth, the registered grey matter images were Jacobian scaled, spatially normalized to the MNI template with a voxel size of 1.5 mm, and then smoothed with a kernel of 4 mm FWHM. Finally, a group-level general linear model was constructed on the grey matter images, with navigation performance as a covariate. To control for potential confounding effects of age and sex, these two factors were included in the model as covariates. To control for different head sizes, total intra-cranial volume was used in global normalization as a covariate.

We created an anatomical mask as our region of interest, which consisted of HPC and ERC in both hemispheres. For each participant, the manually segmented anatomical masks of HPC and ERC in both hemispheres were combined and then normalized to the MNI template. Then, a group-mean mask was obtained by averaging the normalized masks of individual participants. Voxel values in the group-mean mask were between 0 and 1. Finally, the group-mean mask was thresholded by a value of 0; that is, a given voxel was included in the final group mask if it was within the anatomical mask of at least one participant. For voxels in our region of interest, small volume correction approach was adopted to control for multiple comparisons at p < 0.05. For voxels outside of our region of interest, we corrected for multiple comparisons in the entire brain at p < 0.05. Nonparametric permutation tests on linear regression were conducted, using the SnPM toolbox as implemented in SPM12 (Nichols & Holmes, 2002). Covariate of interest was the standardized performance measurement (i.e., landmark-distance error in the landmark condition and walking-direction error in the self-motion condition). Age, sex were entered into the regression model as covariates of no interest. Head size was entered as global value, as a nuisance effect (ANCOVA). Note that the setup of the regression model in VBM analysis was similar to the multivariate linear regression analyses conducted in the ROI volumetry analyses.

For voxel-wise analyses such as VBM, statistical thresholds can be very strict due to the need to control for multiple comparisons. VBM analyses for individual experiments can suffer from inadequate power due to relatively small sample sizes (= 22 participants in each experiment). Therefore, VBM analysis was only conducted on the pooled data, with 44 participants in total. While such an analysis might lose the sensitivity to detect effects specific to individual experiments, it granted us with adequate power to detect true effects common in both experiments.

As shown in Figure 9a, the anterior portion of the left HPC negatively predicted landmark-distance error in the landmark condition (peak t = 4.720, p_FWE-corrected_ = 0.018, MNI coordinate [−21, −15, −12]). Notably, this peak voxel represents the highest significance level across the entire brain. In Figure 9b, partial regression plot was constructed for the mean VBM gray matter of a 4mm-radius spherical area centered at the peak voxel and standardized landmark-distance error, after both variables were adjusted for age, sex, and head size. Landmark-distance error decreased as VBM gray matter estimate increased, and this relationship existed in both individual experiments and the pooled data. No significant effects were observed for positive correlations within the region of interest. For voxels outside the region of interest, no significant effects were observed for either negative or positive correlations. For walking-distance error in the self-motion condition, no significant effects were observed either within or outside the region of interest.

**Figure 9:**
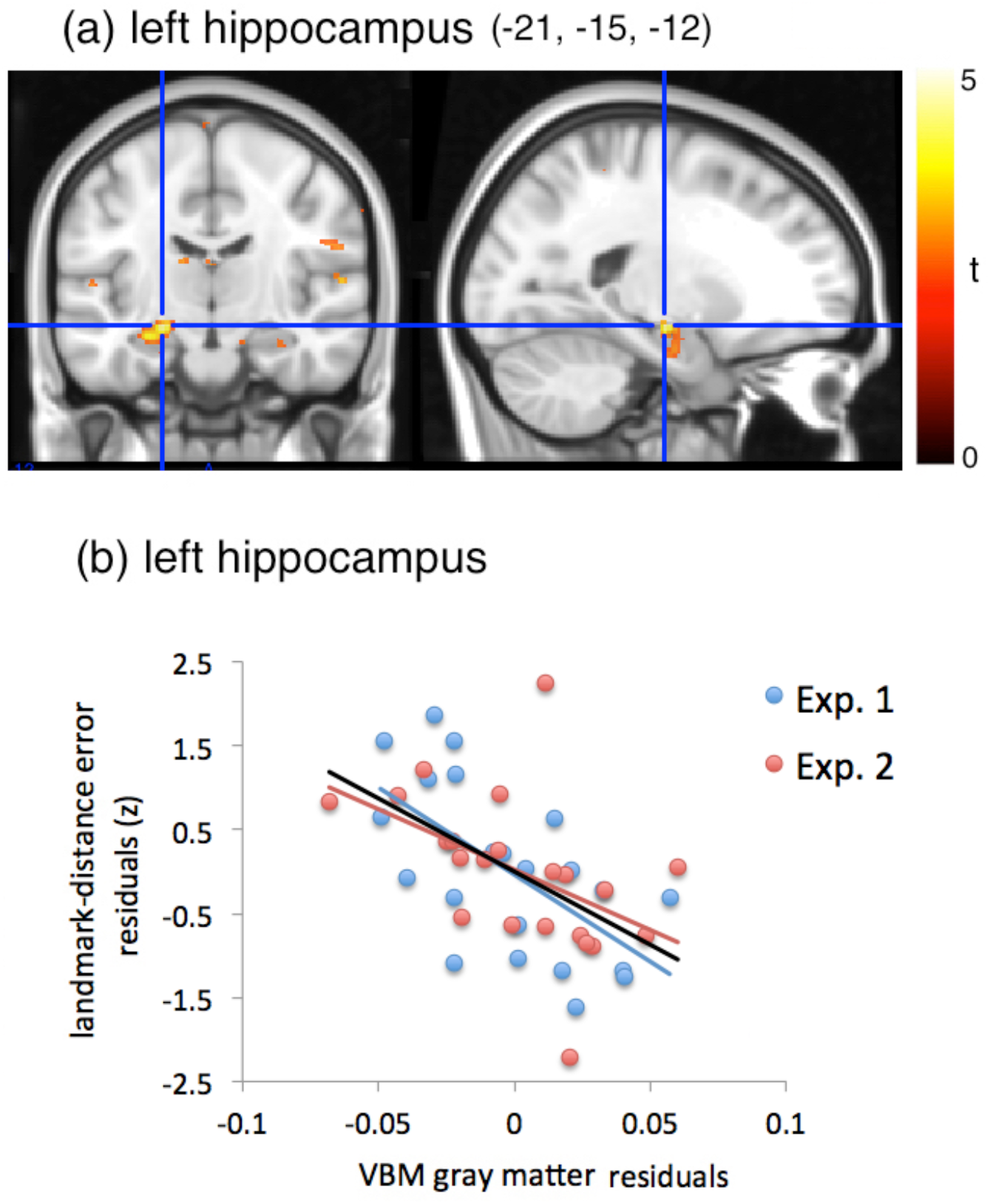
Results of the VBM analysis on the pooled data. (a), the results are displayed on the MNI template at a threshold of p & 0.01 (uncorrected), with the MNI coordinate of the peak voxel indicated. Using the MarsBaR toolbox (Brett, Anton, Valabregue, & Poline, 2002), we created a sphere of 4 mm in radius centered at the peak voxel, and then extracted estimate of the mean gray matter volume for this area. (b), in the partial regression plot, standardized landmark-distance error (z-score) is plotted against VBM gay matter estimate, both adjusted for age, sex, and head size. Data from different experiments are represented with symbols of different colors (blue, experiment 1; red, experiment 2). The linear regression fit lines were shown for the two experiments separately (blue line and red line) and for the pooled data (the dark line).

### 4.3 Discussion

Our analyses of the pooled data again revealed a hippocampal contribution to landmark navigation performance. This effect was localized in the anterior portion of HPC. The highly similar findings in the landmark condition across the two experiments indicate that landmark navigation recruits similar neural mechanisms in both immersive and desktop virtual reality environments. However, unlike Experiment 1, hippocampal volume did not predict performance in the self-motion condition. In addition, unlike Experiment 1, entorhinal volume negatively predicted path integration performance, meaning the larger the ERC, the worse the performance, and this negative correlation was mainly driven by data in Experiment 2 when path integration was based on optic flow but not body-based cues. These discrepancies suggest that path integration was affected by the specific virtual reality setup, and that the hippocampal involvement in path integration might be stronger when body-based cues were used.

## 5 GENERAL DISCUSSION

In our study, participants completed spatial navigation tasks in either a fully immersive virtual reality environment or a desktop virtual reality environment, with landmark and self-motion cues dissociated. The results showed that hippocampal volume consistently and positively predicted navigation performance using a single local landmark in both immersive and desktop virtual environments, and the VBM analysis revealed that this effect was localized in the anterior portion of HPC. We also found that hippocampal volume positively predicted path integration performance in the immersive virtual environment when body-based cues were available. Finally, entorhinal volume was not related to landmark navigation performance, and negatively predicted path integration performance based on optic flow information. Taken together, our study is the first to demonstrate correlations between hippocampal volume and individual differences in navigation performance based on a single local landmark.

Despite different experimental setups and navigation tasks, results of both experiments showed that hippocampal volume positively predicted landmark navigation performance, i.e., the representation of a target’s distance to the local landmark. This suggests that landmark navigation recruits similar neural mechanisms regardless of the specific virtual reality setup. While consistent with previous work showing that hippocampus contributes to processing of landmarks (Bohbot et al., 2007; Hartley & Harlow, 2012; Iaria et al., 2008; Maguire et al., 2000; Woollett & Maguire, 2011), our results demonstrated this relationship when contributions of path integration had been excluded from the navigation performance.

In our study, participants relied on a single local object as the landmark in a given trial in the landmark condition. The local object was a local landmark, as opposed to distal landmarks, since it provided distance information in addition to orientation information (Lew, 2011). Furthermore, participants approached the target location from the same perspective when executing the response. We speculated that egocentric, but not allocentric, representation of the target location was formed and utilized for response, since a view-based matching strategy sufficed to solve the task (Collett & Collett, 2002). Supporting this assumption, a majority of participants reported that during the response stage, they tried to match the size of the landmark as remembered when the landmark was displayed. Relative object size is an important and effective pictorial cue for distance perception (Gillam, 1995; Sedgwick, 1986). Our finding that hippocampal volume predicted behavioural accuracy of retrieving the target’s distance to the landmark suggests that HPC is related to distance estimation based on the view-based matching strategy and therefore contributes to egocentric coding of distance to a single local landmark.

The local landmark used in the landmark condition in our experiments is also classified as a featural cue, as opposed to geometric cues, like a circular enclosure, and configural cues, like multiple landmarks forming an implicit geometric shape. Many studies suggest that – compared to geometric and configural cues – the processing of a local landmark in spatial navigation is not dependent on HPC, but rather on the dorsal striatum (Doeller et al., 2008; McDonald & White, 1994; Packard & McGaugh, 1992, 1996). HPC has been shown to process geometric cues specifically (Doeller et al., 2008; O’Keefe & Burgess, 1996; Wegman et al., 2014) and hypothesized to specialize in allocentric representations of spatial memory (Burgess, 2006; Ekstrom, Arnold, & Iaria, 2014; O’keefe & Nadel, 1978). In contrast to this hypothesis and previous findings, our study suggests that hippocampus also contributes to egocentric encoding of a single local landmark, by demonstrating that hippocampal volume positively predicted navigation performance based on a sole local landmark.

To understand the neural mechanisms of local-landmark navigation, it is critical to clarify how a local landmark can be utilized. First, a single local landmark can be used as an associative cue, such that it is positioned at the reward location or linked to some motor responses (e.g., turning directions) along a route leading to the destination. The associative use of a single local landmark necessitates no extraction of fine-grained spatial information and was found related to the dorsal striatum but not HPC (Hartley, Maguire, Spiers, & Burgess, 2003; Marchette, Bakker, & Shelton, 2011; Packard & McGaugh, 1992, 1996). Second, a single local landmark can be used for precise localization. Our findings suggest that when navigators were capable of extracting fine-grained spatial information from the local landmark, even in an egocentric reference frame, HPC was involved in processing the landmark.

Despite the abundant evidence that place cells are not spatially modulated by local landmarks (Cressant et al., 1997; Cressant, Muller, & Poucet, 1999; O’keefe & Nadel, 1978), a recent rodent study found that firing fields of some hippocampal place cells recorded in the dorsal HPC (homolog to posterior human HPC) lied at fixed angles and distances to local objects in an allocentric reference frame (Deshmukh & Knierim, 2013). This finding leads to the speculation that activities of hippocampal place cells can be utilized to extract fine-grained angular and distance information from local landmarks under certain circumstances. Our finding that the gray matter volume of the anterior HPC predicted distance-to-landmark accuracy with a local object supports the distance proposition of this speculation and suggests that it might be possible to observe similar place cell firings in an egocentric manner in relation to local objects in the ventral portion of the rodent HPC (homolog to the human anterior HPC). Our results did not seem to support the angular proposition of this speculation, since we did not observe significant correlations between hippocampal volume and landmark-angle accuracy in responses.

Previous human fMRI studies pitted a local landmark against a geometric cue - a circular boundary (Doeller et al., 2008), or a configural cue – an object layout (Wegman et al., 2014). Their behavioral measurements tapped on fine-grained spatial information. Both studies showed that HPC was involved in geometric and configural processing, but not related to the processing of the local landmark, inconsistent with our observations. However, in their data analysis, geometric trials and local object trials were contrasted with each other but not with baseline trials, so it was possible that the processing of local objects still involved hippocampal processing, but to a lesser degree compared to geometric cues. This possibility is consistent with the notion that behaviorally, geometric cues and local landmarks might differ long a single dimension of cue salience (Mou & Zhou, 2013). Indeed, a subsequent study by Doeller and colleauges found that hippocampal damage impaired performance with both the local landmark and the boundary (Guderian et al., 2015), in the same navigation task used before (Doeller et al., 2008), implying that hippocampal activity might be related to local-landmark-processing as well.

In addition, Doeller et al. (2008) and Wegman et al. (2014) observed recruitment of the posterior HPC when the geometric and configural cues were used, whereas our study indicated an involvement of the anterior HPC in processing a local landmark, as shown in the VBM analysis on the pooled data. Therefore, our results do not necessarily conflict with these two studies, but rather suggest that different hippocampal portions might support different spatial functions: the posterior HPC supports allocentric representations of the spatial environment, which is usually induced by the use of a geometric boundary (O’Keefe & Burgess, 1996) or an object layout (Mou & McNamara, 2002; Mou, McNamara, Rump, & Xiao, 2006), whereas the anterior HPC is related to egocentric representations, e.g., representing one’s egocentric distance to a local object, as demonstrated in the current study. Consistent with this speculation, previous work has shown that activity in the anterior hippocampus keeps track of the human navigator’s egocentric distance to the goal location (Howard et al., 2014). Our finding is also consistent with the neurophysiological evidence that the perirhinal cortex, which processes object information, projects more to the anterior than the posterior portion of HPC (Libby, Ekstrom, Ragland, & Ranganath, 2012; Maass, Berron, Libby, Ranganath, & Düzel, 2015).

We also found that when body-based cues were involved, hippocampal volume positively predicted path integration performance, i.e., the walking distance to the target location from a fixed location. This relationship was not observed when path integration was based on optic flow inputs. Considering that previous studies using desktop virtual reality setups observed an involvement of the hippocampus in path integration when only optic flow information was provided (Chrastil et al., 2017, 2015; Wolbers et al., 2007), our study suggest that the relationship between HPC and path integration might be stronger when body-based cues were involved. Our findings also suggest that HPC was prominently involved in processing the distance component but not the angular component of the homing vector in path integration. This observation might help explain controversies in lesions studies (Alyan & McNaughton, 1999; Kim, Sapiurka, Clark, & Squire, 2013; Shrager, Kirwan, & Squire, 2008; Whishaw & Maaswinkel, 1998), by proposing the possibility that these inconsistencies might stem from differential involvement of angular vs. distance components in the behavioral measurement.

Finally, we did not observe positive correlations between navigation performance and volumetric measures of ERC. Rather, we observed negative correlations between path integration performance based on optic flow information and ERC volume in Experiment 2. These findings are inconsistent with the abundant evidence from rodent studies that suggest a prominent role of the entorhinal grid cell system in spatial navigation (Moser et al., 2008). However, a recent fMRI study on the grid cell network did not observe correlations between entorhinal BOLD signals and path integration performance in healthy young adults (Stangl et al., 2018). Given the complexity of navigational behavior, we suggest that computations beyond ERC, i.e. in HPC, might be more important for inter-individual differences.

To conclude, we investigated the anatomical basis of landmark navigation and path integration using an individual differences approach. By contrasting path integration and landmark navigation in the same spatial context, our results provide novel evidence implicating HPC in egocentric encoding of a single local landmark and demonstrate a role of HPC in path integration based on body-based cues.

## Acknowledgements

This research was supported by research grant from the Human Frontiers Science Program (RGP 0062/2014, awarded to T.W.)

